# eSVD-DE: Cohort-wide differential expression in single-cell RNA-seq data using exponential-family embeddings

**DOI:** 10.1101/2023.11.22.568369

**Authors:** Kevin Z. Lin, Yixuan Qiu, Kathryn Roeder

**Affiliations:** Department of Biostatistics, University of Washington, Seattle, Washington, United States of America; School of Statistics & Management, Shanghai University of Finance and Economics, Shanghai,People’s Republic of China; Department of Statistics & Data Science, Carnegie Mellon University, Pittsburgh, Pennsylvania, United States of America

**Keywords:** case-control subjects, Gamma-Poisson distribution, matrix factorization, multi-individual data

## Abstract

**Background:** Single-cell RNA-sequencing (scRNA) datasets are becoming increasingly popular in clinical and cohort studies, but there is a lack of methods to investigate differentially expressed (DE) genes among such datasets with numerous individuals. While numerous methods exist to find DE genes for scRNA data from limited individuals, differential-expression testing for large cohorts of case and control individuals using scRNA data poses unique challenges due to substantial effects of human variation, i.e., individual-level confounding covariates that are difficult to account for in the presence of sparsely-observed genes.

**Results:** We develop the eSVD-DE, a matrix factorization that pools information across genes and removes confounding covariate effects, followed by a novel two-sample test in mean expression between case and control individuals. In general, differential testing after dimension reduction yields an inflation of Type-1 errors. However, we overcome this by testing for differences between the case and control individuals’ posterior mean distributions via a hierarchical model. In previously published datasets of various biological systems, eSVD-DE has more accuracy and power compared to other DE methods typically repurposed for analyzing cohort-wide differential expression.

**Conclusions:** eSVD-DE proposes a novel and powerful way to test for DE genes among cohorts after performing a dimension reduction. Accurate identification of differential expression on the individual level, instead of the cell level, is important for linking scRNA-seq studies to our understanding of the human population.

## Background

High-throughput single-cell RNA-seq (scRNA) technology has advanced tremendously over the last decade and helped biologists uncover differing cell-type proportions as well as differentially expressed (DE) genes within a particular cell-type when studying various diseases or disorders. These findings were previously inaccessible using bulk RNA-seq technology. As the technology has developed more in accuracy and cost-efficiency, many labs have begun sequencing entire cohorts of individuals to study up-regulated/down-regulated pathways specific to each cell type in diseases/disorders that generalize to the human population. For example, biologists have sequenced hundreds of thousands of cells for 44 individuals with lung adenocarcinoma [1], hundreds of thousands of neurons for 84 individuals with varying severity of Alzheimer’s Disease [2], and millions of blood cells for 162 individuals with systemic lupus erythematosus and 99 control individuals [3] in order to discover DE genes for the disease/disorder consistent with the entire cohort. While numerous existing and benchmarked single-cell DE methods are designed to find differentially-expressed patterns among *cells* [4, 5], this task fundamentally differs from the cohort-wide studies’ goal of finding differentially expressed patterns among *individuals*. This pressing methodological gap has been raised by many biologists who have collected cohort-wide scRNA-seq datasets [6, 7]. Hence, we reinvestigate the shortcomings of existing DE methods commonly used to analyze cohort-wide scRNA-seq data and design a new DE method specifically suited for such data. Focusing on scRNA-seq data enables us to study more principled ways to model cohort-wide scRNA-seq data, which we hope could be used to inspire cohort-wide DE methods for other single-cell technologies in the future, as well as pioneer applications in future eQTL, as well as upcoming studies that sequence spatial transcriptomics or paired multiomics on large cohorts.

One of the main distinctions between a DE analysis among cells compared to a DE analysis among individuals lies in how the case and control population distributions are quantified. DE analyses among individuals need to account for variability within and among individuals in order to properly model human variation. However, most existing DE methods lack one of these two aspects. On the one hand, variability within an individual hinders “pseudobulk” analyses where all the cells among each individual are summed to yield a pseudobulk sample. Then, methods originally designed for bulk RNA-seq data such as DESeq2 [8] and edgeR [9] are used, but these methods do not account for the variability within an individual. On the other hand, variability across individuals hinders most DE methods made for scRNA-seq data. Specifically, a gene could be differentially expressed among cells but not among individuals if all cells with a significantly higher gene expression come from a small subset of individuals.

The second main distinction is that in cohort-wide scRNA-seq data, there are potentially substantial effects of individual-level covariates such as age, sex, and smoking status that can induce differences in gene expression among the cells that do not reflect the biological differences related to the disease or disorder. A conventional strategy is to regress out the covariate effects for each gene one at a time via a mixed effects model such as MAST [10] and NEBULA [11] where there is a random effect for each individual. However, this regression might be inaccurate since each gene is sparsely sequenced, detrimentally impacting the downstream DE analysis. Hence, an alternative strategy is to use a dimension-reduction method via matrix factorization such as GLM-PCA [12], ZINB-WaVE [13], or scGBM [14]. These methods pool information across genes to remove confounding covariates’ effects more effectively. However, a naive application of DE testing on the dimension-reduced scRNA-seq data has been observed to inflate the Type-1 error [15]. This inflation occurs because the dimension reduction introduces correlations among genes that contaminate the signal. Null genes could seem significantly (and erroneously) differentially expressed after a dimension reduction if the test is not performed with care. (See an illustration of this phenomenon in Supplementary Figure A1.) The main focus of our paper is to solve this statistical dilemma of how to perform differential testing after dimension reduction. We note that another recent line of work has developed differential testing among cells after a dimension reduction via deep variational autoencoders [16, 17]. The authors overcome the aforementioned pitfall of DE testing after dimension reduction by leveraging the inherent randomness in autoencoders and adding pseudocounts to avoid deeming genes with a small log-fold change as significant. However, we do not pursue this direction since matrix factorizations offer practitioners a more transparent and interpretable framework.

We design the exponential-family SVD differential expression (eSVD-DE) to overcome these two main obstacles, which extends our previous work [18]. Importantly, our method infers the differential expression based on the posterior distribution after performing a dimension reduction, which helps counteract the Type-1 error inflation. This combination of matrix factorization, the posterior distribution, and a test statistic designed to assess differential expression among individuals enables eSVD-DE to better detect DE genes in cohort studies compared to current methods. The eSVD-DE also enables model diagnostics to assess if the assumed statistical model is appropriate for modeling the scRNA-seq dataset.

In this paper, we focus on testing for cells of a particular cell type. We show that eSVD-DE can find reproducible signals in multiple pairs of cohort datasets, either across various cell types between two independent studies of idiopathic pulmonary fibrosis (IPF) in the human lung [6, 19] or within a study of non-inflamed and inflamed cells studying ulcerative colitis in the human colon [20]. We also show that eSVD-DE can find novel DE genes across different cell types in a dataset studying autism [21]. Altogether, these analyses provide evidence that eSVD-DE is a valuable tool for investigating differential expression among cohort studies that will become more prevalent as high-throughput sequencing technologies are applied to large cohorts.

## Results

### Overview of eSVD-DE for differential expression testing

eSVD-DE performs DE by first projecting the cells onto a low-dimensional manifold while removing the effects of covariates. This step is the cornerstone and namesake of our method. Our dimension reduction follows previous dimension-reduction work such as GLM-PCA [12], ZINB-WaVE [13], and scGBM [14], where we embed the cells via the Poisson distribution based on the gene expression *A* ∈ {0, 1, 2, … } ^*p×n*^ and the covariate matrix *C* ∈ ℝ^*n×r*^, where *n, p*, and *r* denote the number of cells, genes, and covariates (Figure 1A). Importantly, these covariates contain an intercept term, the log sequencing depth (computed as the log of the total counts per cell), the case-control indicator, and covariates that could be potential confounders, such as clinical covariates of each individual (sex, age, and smoking status). We denote the

**Fig. 1:**
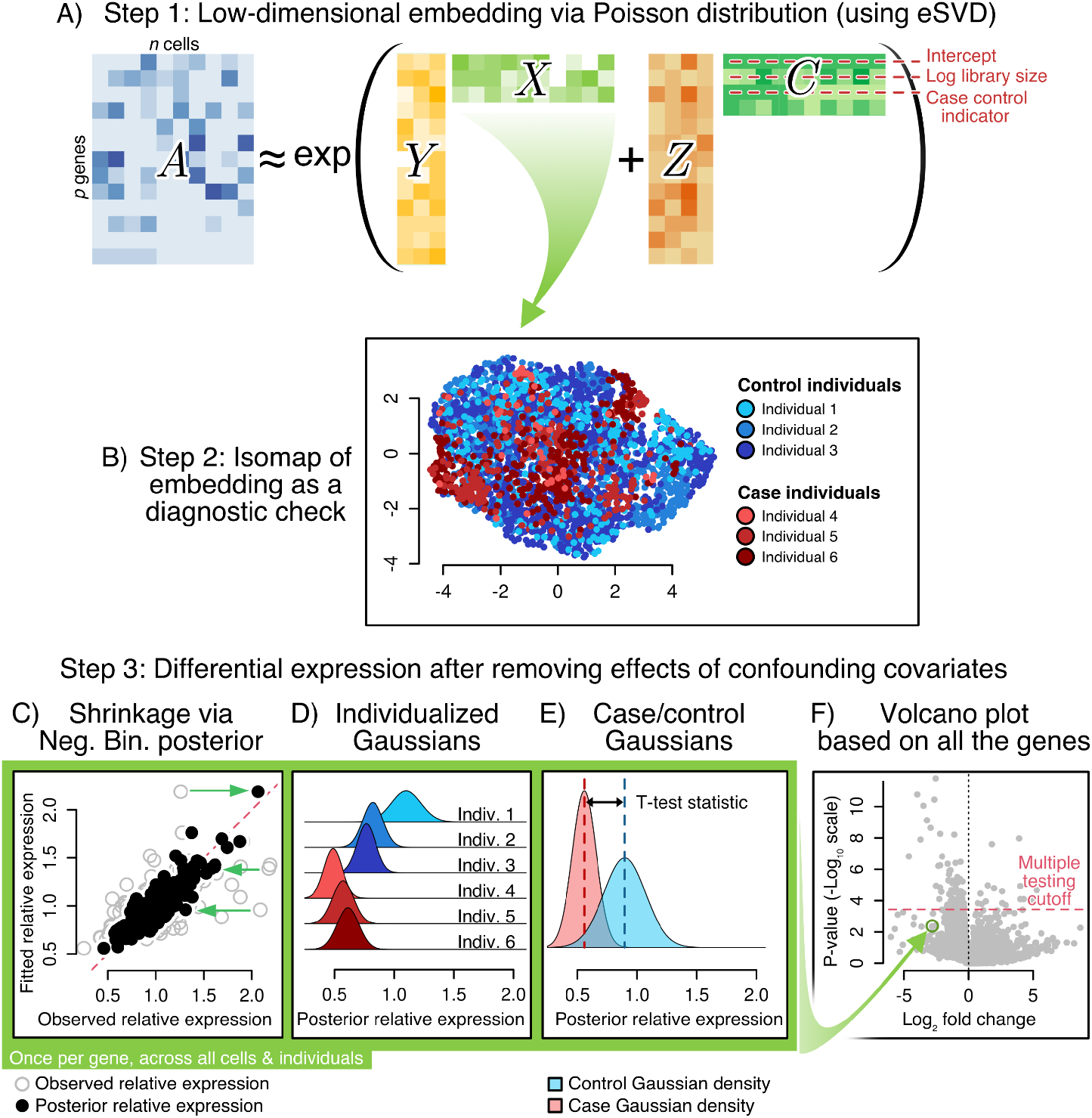
**A** Schematic of the eSVD-DE’s matrix factorization, where the observed scRNA-seq data is modeled as a sum of two low-rank matrices, one for the covariates and one for the cells’ latent vectors, with an exponential link function (for the Poisson distribution). **B** The cells’ latent vectors can be used for diagnostic checks, such as visualization via Isomap. **C** To account for overdispersion and over-fitting of the dimension reduction, shrink each cell via the negative binomial distribution’s posterior mean. **D** Represent each individual by a Gaussian distribution among the individual’s cells. **E** Compute a test statistic analogous to the T-test after aggregating cells from the cases or control individuals in the cohort. C through E are performed for each gene. **F** Volcano plot, showing a multiple testing cutoff to determine the significant DE genes.

specific covariate for the case-control indicator as *C*_.,(cc)_. The eSVD-DE learns a coefficient matrix that removes the effects of the covariates as well as the low-dimensional embedding of “residuals” *X* via the hierarchical model for gene *j* ∈ {1, …, *p*} and cell *i* ∈ {1, …, *n*} following work such as [25],

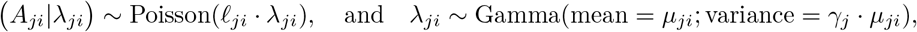

and

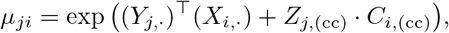

where *X* ∈ ℝ^*n×k*^ and *Y* ∈ ℝ^*p×k*^, and 𝓁_*ji*_ denotes the covariate-adjusted sequencing depth that accounts for the effect of all the remaining covariates in *C* aside from *C*_.,(cc)_. Here, *γ*_*j*_ denotes the overdispersion parameter for gene *j*, which measures how much extra variability exists between the assumed Poisson fit in the low-dimensional embedding and the observed data, analogous to works like SAVER [22] and totalVI [23]. While this *k*-dimensional embedding *X* can be helpful for a wide variety of applications downstream, we focus on modifying our previous work [18] to perform DE analysis for cohort data. We advocate using a matrix-factorization-based approach to perform this dimension reduction over deep learning approaches since our approach enables conventional model diagnostics to assess and tune eSVD-DE (Figure 1B).

After fitting a low-dimensional embedding, we cater the following procedure to test for DE among individuals, an aspect absent from our previous work [18]. First, we adjust the predicted relative expression for each cell’s gene expression using its posterior according to the Gamma-Poisson distribution (i.e., Negative Binomial, Figure 1C). This posterior, denoted by *λ*_*ji*_ | *A*_*ji*_, is computed by first estimating the overdis-persion *γ*_*j*_ of each gene. Importantly, this posterior distribution is computed to be the relative expression after accounting for the contributions of the confounding covariates. For a particular gene, we summarize the cells’ posterior distribution of *λ*_*ji*_ | *A*_*ji*_ from each individual as a Gaussian distribution following the Central Limit Theorem, and we then compute the T-test statistic reflecting the collective difference between the Gaussians from case individuals to those from control individuals (Figure 1D-E). Notably, this means we do *not* perform a differential expression test based on *µ*_*ji*_’s because different genes are highly correlated based on their values in *µ*_*ji*_ due to the low dimensional embedding, which will artificially inflate the Type-1 error. (See the Appendix for a more in-depth discussion.) Finally, we use a multiple-testing procedure based on empirical null distribution to report the DE genes, which has been successful in other settings to account for possible model misspecification [24] (Figure 1F). More details of the eSVD-DE procedure are described in “Statistical model and method.”

eSVD-DE relies on three primary statistical assumptions: (1) scRNA-seq data can be appropriately modeled through the Gamma-Poisson distribution, (2) the effects of confounding covariates can be effectively removed through a GLM framework, and (3) the DE genes show significant differences in means between case and control individuals after accounting for the individual-level covariates. Towards the first assumption, our posterior distribution effectively models counts through a Negative Binomial distribution, which has been justified for modeling scRNA-seq data [25] and has served as the foundation for many methods [13, 22, 26]. Towards the second assumption, many methods from the bulk RNA-seq data such as edgeR [9] and DESeq2 [8] have found tremendous success regressing out covariates through the GLM framework. Towards the last assumption, we note that there are existing methods such as IDEAS [27], scDD [28], and waddR [29] that test for differential *distributions* instead of differential mean expressions. However, we have found it more challenging to generalize these results when comparing different datasets of similar biological systems, as differentially distributed genes are not neatly characterized by over-/under-expression.

### Design of simulation studies

The main message we wish to convey in this simulation section is two-fold: 1) testing for differential expression among individuals is fundamentally different from among cells, and 2) eSVD-DE’s framework enables more accurate inference due to its usage of dimension reduction and posterior correction. Figure 2A conceptualizes the first point. Large within-individual variability hinders “pseudobulk” DE methods that sum the gene expressions across all the cells originating from the same individuals since these methods do not account for the variability of expression within an individual. On the other hand, large between-individual variability hinders most existing methods designed for testing DE genes among cells from scRNA-seq data. This is because even if the difference in mean expression is insignificant on the cohort level, many cells from a small subset of individuals can yield highly significant differences in mean expression on the cell level. Since previous work has explained the importance of accounting for within-individual variability [27, 30, 31], we focus on a simulation that illustrates the importance of accounting for the between-individual variability here.

**Fig. 2:**
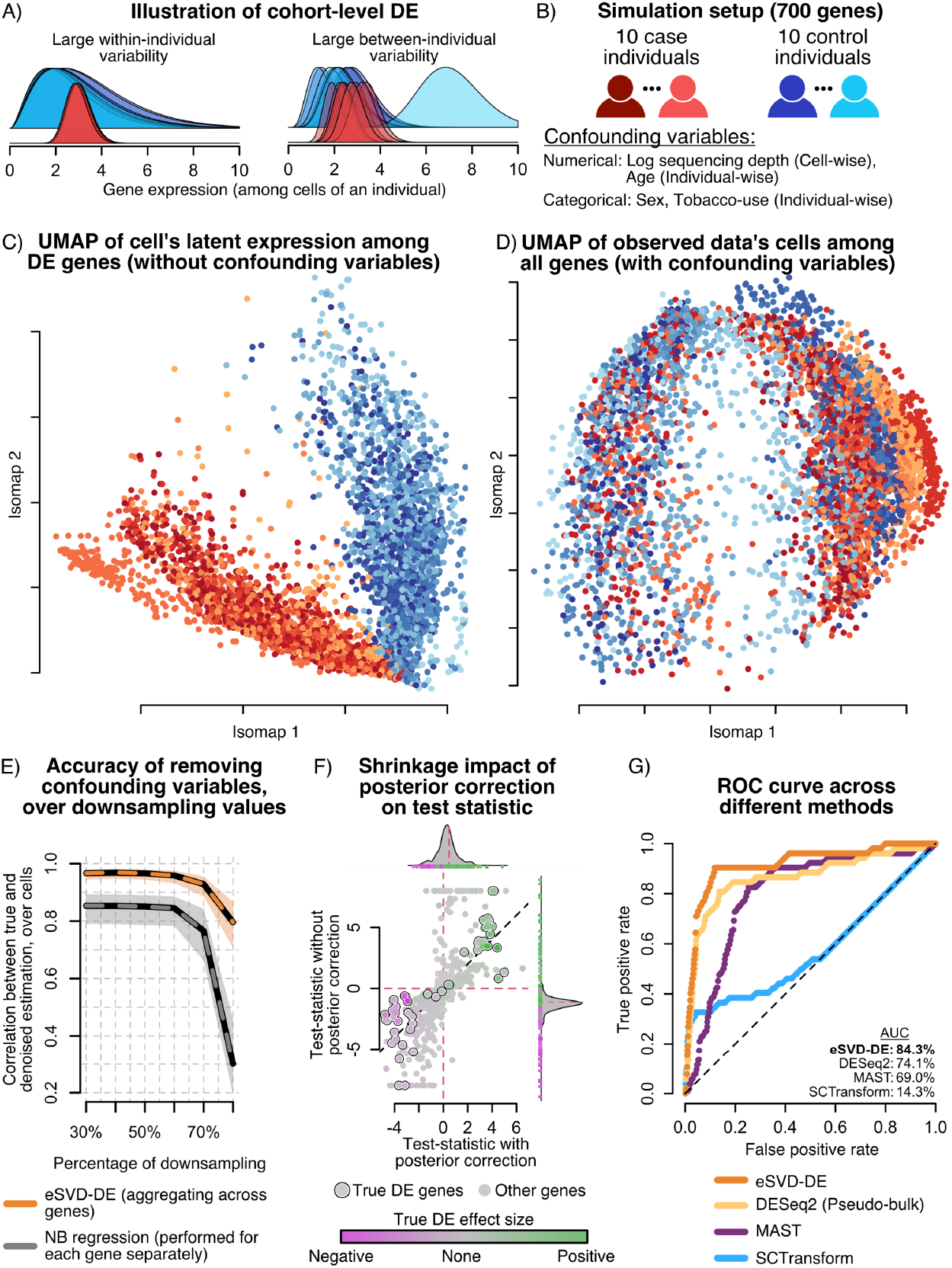
**A** Illustration of challenges for cohort-wide DE testing. **B** Setup for our simulation setup. **C** Isomap of the cells based on the true DE genes’ expression before introducing the confounding variables. No individuals concentrate tightly in any region on the Isomap manifold, and there is a strong separation between the cases (shades of red) and controls (shades of blue). **D** Isomap of the observed data based on all the genes. Cells from the same individual concentrate in the embedding, suggesting that confounding covariates additionally drive the difference in expression profiles among individuals. **E** Downsampling experiment, demonstrating that by pooling information across genes, eSVD-DE outperforms gene-by-gene Negative Binomial regression for regressing out covariate effects. **F** Illustration of the importance of shrinkage, where the x-axis and y-axis represent each gene’s test statistic with and without posterior correction, respectively. The genes are colored by their true log-fold change, of which the circled genes denote the top 50 genes with the highest true log-fold change. **G** ROC curve comparing four different methods, illustrating that eSVD-DE has more power than competing methods. The area under the curve (AUC) is shown for each method, where the percentage represents the area between the method’s curve and the diagonal line as a fraction of total possible area. The bolded method denotes the method with highest AUC.

Our simulation study contains 700 genes and 20 individuals equally split among cases and controls, where each individual contributes 250 cells. The cells have gene expression profiles that are impacted by covariates correlated with the case-control status (such as age, sex, and tobacco use) and are sampled from a Gamma-Poisson distribution (Figure 2B). Overall, the true generative model is more complex than the statistical model assumed by eSVD-DE to not give eSVD-DE an unfair advantage in our benchmarking experiment. Nonetheless, the data is generated such that 50 genes are drastically more differentially expressed among cases and controls on the cohort level than the remaining 650 genes, barring confounding effects. This true signal can be visualized via an Isomap [32, 33], where there is no apparent stratification within the case or control individuals (Figure 2C). However, when the covariate effects of age, sex, and tobacco use are also included, there is obvious confounding of which genes are differentially expressed (Figure 2D). Specifically, the cells from the same individual concentrate in different regions of the embedding. We choose to use the Isomap here, as opposed to the more commonly used UMAP [34], since the Isomap has well-studied statistical properties [35–37]. This quality is valuable when we use these visualizations to diagnose our hypothesis testing framework instead of an exploratory tool, as we will illustrate later.

### Simulations verify that eSVD-DE effectively remove covariate effects in the presence of sparsity

We first show that by pooling information across genes via the dimension-reduction framework, eSVD-DE removes the confounding variables’ effects more accurately compared to a Negative Binomial (NB) regression applied separately for each gene (Figure 2E). To demonstrate this, we incrementally downsample the data to reduce the signal size of the DE genes. As the amount of downsampling increases, it will be more difficult for a method to remove the confounding effects properly. We measure the accuracy of how well the confounding effects were removed by computing the Pearson correlation between each cell’s vector of true natural parameters across the 700 genes and its vector of estimated natural parameters after the confounding effects have been removed. We observe that at starting at 30% downsampling, eSVD-DE removes the confounding effects more accurately than the gene-by-gene NB regression (0.95 to 0.87). Additionally, although both methods drop in accuracy for downsampling levels of 60% or more, eSVD-DE is still more accurate than the gene-by-gene NB regression. This finding is statistically intuitive since when genes are sparsely observed, there is not enough information within a specific gene to accurately estimate the NB regression coefficients. However, by pooling information across genes, the coefficients for the confounding variables are better estimated.

### Simulations verify that eSVD-DE’s posterior enables gene-specific discoveries

We next illustrate that the posterior correction performed by eSVD-DE is important for DE testing on the cohort level (Figure 2F). We plot the test statistic derived directly from the low-dimensional embedding against the test statistic derived from the posterior-corrected relative expression, where we mark the 50 genes the largest true log-fold change. In particular, the former represents the prototypical analysis of applying DE after denoising the data by a dimension-reduction method, and we observe that many null genes display large test statistics (x-axis), confirming the findings of previous work that observes an inflation of Type-1 error for such procedures [15, 30, 38]. However, by adjusting the relative expressions using the posterior mean, many null genes have drastically smaller test statistics (y-axis). This phenomenon occurs because the assumed linear model between the covariates and the gene’s natural parameter does not sufficiently capture more complex non-parametric relationships often displayed among individuals, distorting the lower-dimensional embedding of the “residuals.” This distortion has an adverse effect when genes vary substantially in sparsity – sparsely observed genes are denoised by projecting the cells onto the incorrect manifold.

### Simulations verify that eSVD-DE’s low-dimensional embedding improves power over other methods

Lastly, we illustrate the eSVD-DE has more power than other conventional methods to test for DE in cohort-wide scRNA-seq data through our simulation. Since different methods estimate sets of DE genes with dramatically different sizes for a particular FDR cutoff, we plot the entire curve of true positive rate (TPR) and false positive rate (FPR) over all possible cutoffs (Figure 2G). Here, we compare against three other methods: DESeq2 [8] (i.e., the prototypical method representing “pseudobulk” analyses), MAST [10] (i.e., the commonly used method using mixed-effect models where there is a random effect among individuals) and SCTransform [39] (i.e., the prototypical method of performing DE ignoring the individual structure). We observe that eSVD-DE has the highest power compared to the three other methods. DESeq2 performs the best among the three competing methods since the averaging among cells of an individual dramatically reduces the estimation variability, while MAST performs the next best since it accounts for the individual structure.

We perform further power analyses across different number of genes and imbalances between number of cells across individuals (Supplementary Figure C4 and C5). We also perform a separate simulation in the Appendix (Supplementary Figure B2 and B3) to demonstrate that eSVD-DE does not inflate the Type-1 error among true null genes.

### eSVD-DE enables diagnostics to assess the performance of removing covariate effects

Moving beyond simulations, we now investigate how well eSVD-DE removes confounding effects in scRNA-seq datasets and how the dimension-reduction framework enables conventional model diagnostics. To demonstrate this, we investigate a broad collection of scRNA-seq datasets from diverse tissues but focus here on the 8909 T-cells from a dataset of lung cells containing 10 healthy individuals and 24 individuals with IPF sequenced using the 10x Chromium single-cell platform, henceforth called the “Adams” dataset [6]. We focus on 6969 genes for our analysis, including 5000 highly-variable genes, the genes reported to be DE by the authors as well as the 1003 human housekeeping genes [40]. We report the summary table of the Adams dataset, as well as the other datasets in this paper, in Table 1, where *p* denotes the number of genes and *n* denotes the number of cells. We include an extended summary table of the datasets in the Appendix When visualizing these cells via Isomap, we can see a clear separation of the case and control individuals, suggesting there are many genes with separable expression patterns (Figure 3A). However, we also see that individual-level covariates like sex and smoking status also are locally concentrated in different regions of the Isomap. We do not wish to report genes as differentially expressed if the differences are induced only by sex differences or smoking.

**Table 1:**
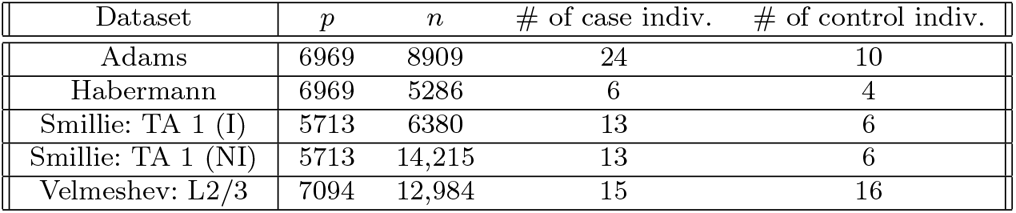
Summary table of datasets.

**Fig. 3:**
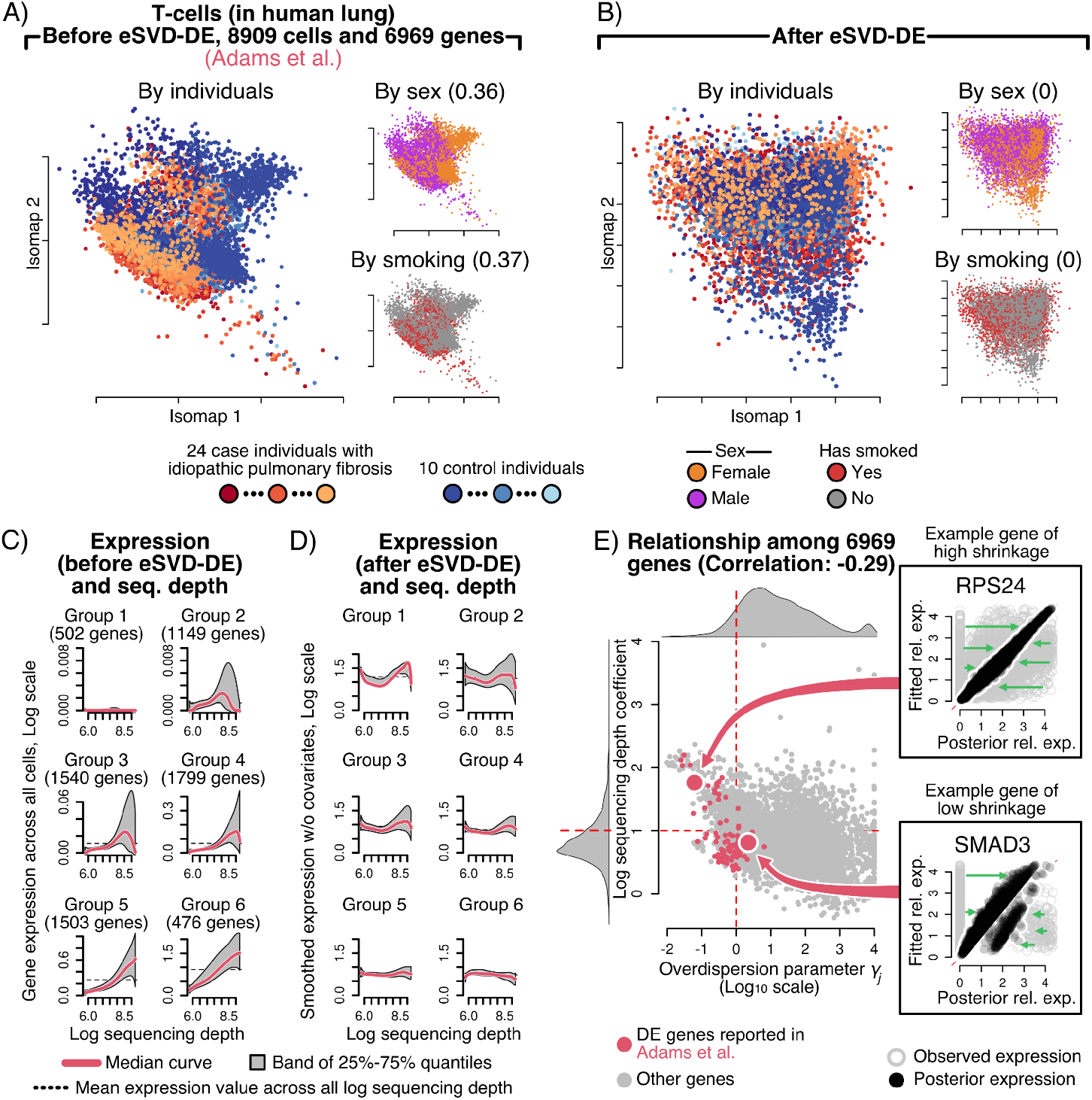
**A** Isomap of the leading principal components for the T-cells in the Adams dataset, shown by individuals, sex, and smoking status. The number in the insets denotes how correlated the individual-level covariate is correlated with the leading principal components. **B** Isomap of the cell embedding after applying eSVD-DE, demonstrating that confounding covariate and individual effects have been removed. The number in the insets denotes how correlated the individual-level covariate is correlated with the estimated eSVD-DE cell embedding. **C**,**D** Relationship between gene expression and cells’ log sequencing depth before or after eSVD-DE (shown on a log scale), respectively, for groups of 6 genes partitioned by the gene’s mean expression relationship after eSVD-DE. After eSVD-DE, each gene becomes more uncorrelated with the cell’s log sequencing depth. **E** Relationship between each cell’s log sequencing depth and overdispersion parameter, highlighting two genes displaying different shrinkage levels via the posterior mean.

After applying eSVD-DE, we see that the resulting Isomap of the fitted embedding no longer carries expression patterns correlated with the individuals, sex, or smoking status (Figure 3B). If biologists prefer quantitative diagnostics, the practitioner can purposely omit a small percentage of values from the scRNA-seq count matrix before applying eSVD-DE and assess how correlated the predicted values are to these omitted values. This strategy was used successfully in our previous work [18]. For all these reasons, we advocate the matrix-factorization strategy in the eSVD-DE as opposed to deep-learning alternatives where there are fewer interpretable diagnostics available. The remaining diagnostics and the Isomaps for other scRNA-seq datasets are shown in the Appendix (Supplementary Figure F6).

### eSVD-DE appropriately adjusts for the sequencing depth using a dimension-reduction approach

Many authors have concluded that appropriately adjusting for the sequencing depth of each cell is one of the most influential aspects of an effective DE method [41, 42]. This adjustment is critical for scRNA-seq data since we do not wish to deem genes as DE simply because specific cells have a larger sequencing depth than others. Instead, genes should be deemed as DE if the case and controls have significantly different expression relative to the cells’ sequencing depth. Accounting for this sequencing depth has been the source of numerous debates on how to normalize the cells’ expression best [43]. The most commonly used approach is to log-normalize the gene expressions, but many existing work has cited that this normalization distorts qualitative aspects of the scRNA-seq dataset [12, 39].

In order to quantify how well the sequencing depth effect was removed from the scRNA-seq data, we perform diagnostics analogous to those in [39]. Specifically, we group the genes into 6 bins based on their mean gene expression. We observe a significant relationship between each gene’s expression and the cells’ sequencing depth before normalizing the scRNA-seq data (Figure 3C). This means without any accounting of the sequencing depth, genes could be deemed significantly differentially expressed solely due to cells having different sequencing depths. However, after fitting the eSVD-DE, this relationship is mainly removed across all bins, discounting the genes with the smallest mean expressions (Figure 3D). Observe that eSVD-DE bears an advantage over other normalization methods such as SCTransform [39] and Scran [44] since eSVD-DE assesses the appropriate sequencing-depth normalization by simultaneously accounting for the cells’ gene expression and covariate information through its dimension-reduction framework, an important quality when dealing with cohort data. See Supplementary Figure F7 for the application of this diagnostic on other datasets, and Supplementary Figure F8 for how this diagnostic performs when only log-normalization is performed instead of the eSVD-DE in the Appendix.

Lastly, recall that our eSVD-DE framework adjusts the fitted gene expression values by the posterior Negative Binomial distribution. This adjustment relies on first 10 quantifying the overdispersion of each gene, i.e., a higher overdispersion means the gene’s expression patterns conform less to the estimated low-dimensional embedding. We hypothesize a negative correlation between the sequencing-depth normalization and the amount of overdispersion. This hypothesis is in line with previous works that investigate this relationship [39], citing that a smaller overdispersion implies that the fitted values are a good approximation of the gene expressions, which often occurs for densely observed genes. This can be visualized as a scatterplot (Figure 3E). Additionally, this plot enables practitioners to survey the amount of shrinkage across all the genes broadly. We highlight RPS24 and SMAD3, 2 genes implicated in previous studies [45–47], as example genes that are highly and lowly shrunk via the posterior distribution since these genes were. All these aspects collectively support the claim that eSVD-DE is appropriately adjusting each gene’s expression by the sequencing depth, which enables us to investigate the performance of the DE analyses downstream.

### eSVD-DE recovers reproducible differences between case and control expression across multiple datasets

While it is difficult to assess the validity of DE genes in cohort-wide scRNA-seq data due to the lack of negative control genes, we hypothesize that a reliable proxy to assess the quality of the DE method is to collect two different cohort-wide scRNA-seq datasets of the same system and diseases/disorder and see if genes reported to be significant in one dataset display similar significances in the other. Towards this end, we investigate another dataset of 5286 T-cells of lung cells from 4 healthy individuals and 6 individuals with IPF sequenced using the 10x Chromium single-cell platform of the same 6969 genes here, henceforth called the “Habermann” dataset (Figure 4A) [19]. Since we are primarily interested in comparing eSVD-DE to other DE methods using this pair of datasets, we do not deploy a multiple-testing procedure to select DE genes but investigate the sets genes with the largest test statistic magnitudes of the same cardinality as those reported by the authors for meaningful comparisons. For starters, the volcano plot on the Adams dataset shows that 20 of the 84 genes with the largest test statistics derived from our eSVD-DE procedure intersect with the 84 DE genes reported by the authors, resulting in Fisher’s exact test p-value of 2.2 *×* 10^−21^ (Figure 4B). Similarly, the volcano plot on the Habermann dataset shows that 30 of the 157 genes with the largest test statistics derived from our eSVD-DE procedure intersect with the 157 DE genes reported by the authors, resulting in Fisher’s exact test p-value of 3.1 *×* 10^−20^ (Figure 4C). Additionally, since housekeeping genes constitute genes primarily responsible for basic cellular functions and are stably expressed regardless of cellular condition, we hypothesize that these genes should not carry substantial differential expression patterns [40]. This phenomenon is demonstrated through the volcano plots (Figure 4B,C).

**Fig. 4:**
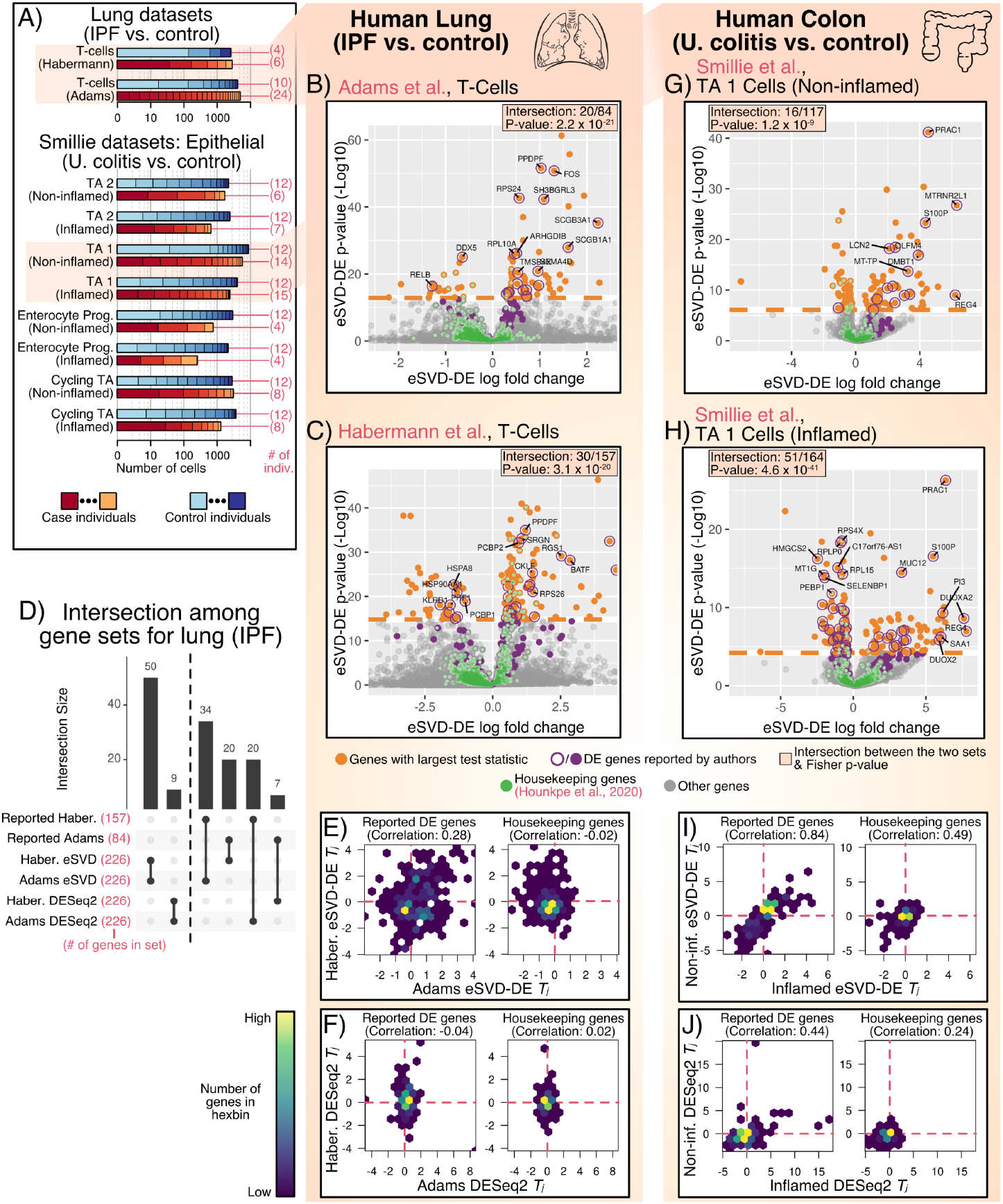
**A** Number of case and control individuals and the total number of cells across cell types and datasets. The x-axis denotes the total number of cells on a log scale, while the partitioning of case and control cells denotes the number of cells from each individual relative to the total number of cells. **B**,**C**,**G**,**H** Volcano plot, where the set of genes with the largest test statistics (having the same size as the original author’s set) is depicted in orange. The genes reported by the respective authors or housekeeping genes are in purple and green, respectively. The Fisher exact test’s p-value between the enrichment of eSVD-DE’s DE genes and the author’s reported DE genes is also reported. **D** Upset plot showing the intersection between pairs of DE genes, either the reported DE genes in the Habermann and Adams dataset and the estimated DE genes using eSVD-DE or DESeq2. **E**,**I** Hexplots showing the correlation between genes’ eSVD-DE test statistics across datasets, either the originally reported DE genes in either dataset or the housekeeping genes. **F**,**J** Similar to **E**,**I** but showing the gene’s DESeq2 test statistics.

When comparing different DE methods, we ask if the leading genes derived from the Adams dataset using eSVD-DE are either (1) genes reported by authors of the Habermann dataset or (2) themselves the leading genes derived from the Habermann dataset using eSVD-DE, or vice-versa. If the size of this intersection is larger than those derived by other DE methods in either scenario, we would have partial evidence that eSVD-DE is more consistently recovering reproducible signals across both datasets. We compared against DESeq2 whereby cells of each individual are aggregated into “pseudobulk” expressions prior to performing DESeq2 [8] for this investigation. We highlight this pseudobulk procedure specifically since this procedure was reported to be more reliable for scRNA-seq with biologically-replicate samples, but its reliability for cohort data was not investigated [4]. For this comparison, we select the 226 genes with the largest test statistics in magnitude from either the Adams or Habermann dataset using either eSVD-DE or DESeq2 since the 84 reported DE genes for the Adams dataset and 157 reported DE genes for the Habermann dataset yield 226 unique genes. The upset plot shows that eSVD-DE yields larger intersections between the two datasets than DESeq2 (Figure 4D).

Additionally, we hypothesize that if a DE method recovers reproducible signals, the DE genes test statistics should be positively correlated between the two datasets. Indeed, if a reported DE gene has a positive log-fold change in one dataset, the other dataset should also show evidence of a positive change. On the other hand, we hypothesize that genes unrelated to the disease/disorder should be uncorrelated test statistics between the two datasets. This hypothesis would be biologically justifiable, as the log-fold change of a gene unrelated to disease/disorder would be determined by random chance. We observe these relationships for eSVD-DE’s test statistics (Figure 4E). In contrast, the correlation for DESeq2’s test statistic among the reported DE genes is near-zero (Figure 4F). We suspect all the above phenomenons are likely driven by pseudobulk methods’ lack of accounting for the within-individual variability.

To further support our empirical claims regarding eSVD-DE, we also study cells from 18 individuals with ulcerative colitis (UC) and 12 healthy individuals across multiple cell types in the colon and 5713 genes, henceforth called the “Smillie” dataset [20]. Each individual with UC contributed cells from inflamed and non-inflamed colon biopsies, and each healthy individual contributed two biologically replicated samples from biopsies in analogous colon regions. In our analysis, we treat each tissue sample as a different “individual” and perform two DE analyses – one of non-inflamed samples from individuals with UC against healthy samples (i.e., the “non-inflamed” analysis) and another of inflamed samples from individuals with UC against healthy samples (i.e., the “inflamed” analysis). Importantly, we split the healthy tissue samples so each healthy sample is only involved in one of the two DE analyses. While we apply this pair of analyses on multiple cell types in this biological system, we focus on the analysis of transit amplifying 1 (TA 1) cells here (Figure 4A). Similar to before, we see that the volcano plots for both the non-inflamed and inflamed DE analysis yield highly-enriched intersections between the genes with the largest test statistics and the DE genes reported by the authors, as well as near-zero enrichment of the housekeeping genes (Figure 4G,H).

The authors of the Smillie dataset observed a high correlation among the test statistics of reported DE genes between the non-inflamed and inflamed DE analyses when aggregating across all the cell types, suggesting that the transcriptomic signature of UC precedes inflammation [20]. We hypothesize that a higher-powered DE analysis of cohort-wide scRNA data should reveal a strong correlation even when focusing on only one cell type. Indeed, we see this positive correlation among eSVD-DE’s test statistics (*ρ* = 0.84, Figure 4I). Additionally, while we see a positive correlation among the housekeeping genes (*ρ* = 0.49), many of such genes have a near-zero test statistic in the non-inflamed analysis. This observation suggests that the differences in housekeeping genes’ expression are induced more by the inflammation of the tissue rather than a biological mechanism disrupted by ulcerative colitis. In contrast, when we apply a similar pair of analyses using DESeq2 on pseudobulk data, the correlation among the reported DE genes is substantially lower (Figure 4J).

### eSVD-DE detects DE genes highly enriched with previously-annotated genes

Having demonstrated all the advantages of eSVD-DE, we hypothesize that our method leads to novel cell-type specific DE discovery. Towards this end, we analyze brain single-cells from controls and case individuals who had been diagnosed with autism spectrum disorder (ASD), partitioned among many cell-types [21], henceforth called the “Velmeshev” dataset (Figure 5A). We focus on this particular system since the Simons Foundation Autism Research Initiative (SFARI) routinely curates a list of autism risk genes based on genetic studies from many publications [48], which can provide relevant genes to compare our eSVD-DE results against. Additionally, a recent bulk-DE analysis of ASD provides a high-quality and independent list of genes to compare against, which we will call the “bulk DE genes” henceforth [49]. A priori, we would not expect any method deployed on the Velmeshev dataset to fully match these lists because our analysis is cell-type specific and the SFARI list are genetic risk genes, which do not always lead to differential expression.

**Fig. 5:**
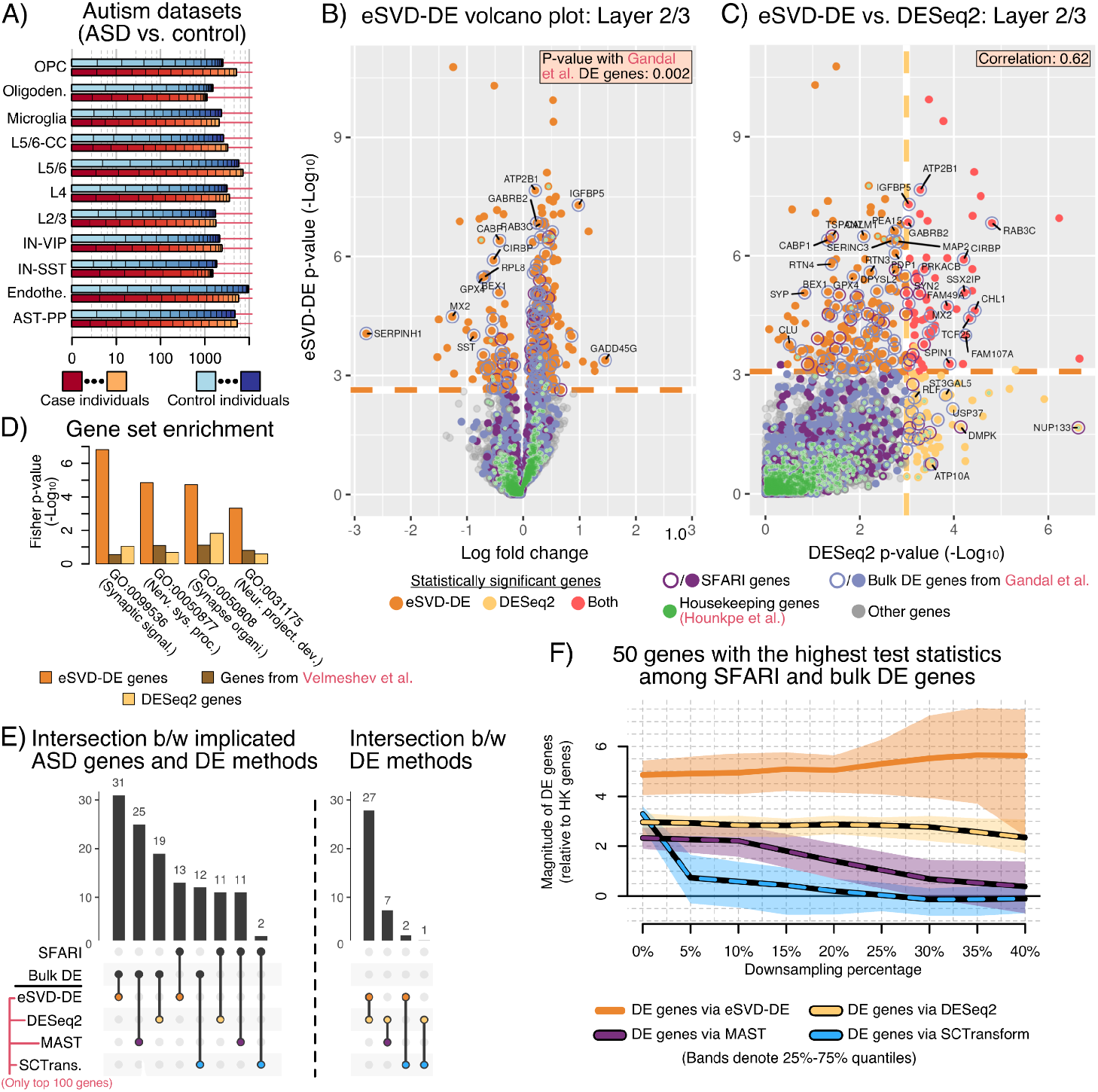
**A** Number of case and control individuals and the total number of cells across cell types for the Velmeshev dataset, analogous to Figure 4**A**. **B** Volcano plot showing eSVD-DE’s results for layer 2/3 cells, where the set of genes exceeding the empirical FDR cutoff are depicted in orange. The SFARI, bulk DE, and housekeeping genes are in purple, blue, and green, respectively. The Fisher exact p-value between the selected genes via eSVD-DE and the bulk DE genes is noted. **C** Comparison of the eSVD-DE p-values against the DESeq2 p-values, and the genes exceeding an FDR cutoff for DESeq2 are shown in yellow. The correlation between the two sets of negative log_10_ p-values is noted. **D** Increase in enrichment for relevant GO terms among the 331 DE genes found by eSVD-DE, the 144 DE genes found by DESeq2, and the 109 genes initially reported by the authors. **E** Upset plot comparing the SFARI or bulk DE genes with the top 100 DE genes estimated by eSVD-DE, DESeq2, MAST, or SCTransform. **F** Downsampling experiment, illustrating the stability of the 50 genes with the largest test statistics in magnitude among the SFARI or the bulk DE genes as the dataset is artificially downsampled. The values on the *y*-axis is the ratio of the test statistic between these genes and the housekeeping genes.

When analyzing the 12,984 cells in layer 2/3, we observe that eSVD-DE estimates 331 DE genes (among the 7055 genes used in the analysis) using an empirical FDR cutoff of 0.05, where we calibrate the p-value according to an empirical null distribution to account for potential misspecification [50] (Figure 5B). Here, 42 and 95 of these genes are among the SFARI and the bulk DE genes, respectively (among the total 800 and 1556 genes, respectively). Comparing the bulk DE genes to our eSVD-DE genes (which are both derived from transcriptomics data), we obtain a significant Fisher p-value of 0.002. Additionally, when we compare DESeq2 for the same cells to find DE genes (144), only 19 and 30 genes overlapped with the SFARI and the bulk DE genes, respectively (Figure 5C). This finding demonstrates that eSVD-DE can find more DE genes, which are more relevant, compared to DESeq2. We report analogous plots in the Appendix when comparing against other methods, such as MAST and SCTransform (Supplementary Figure F9). We note that we do not compare against the DE genes found by the authors of the data themselves here, as they had used MAST to find their DE genes. As our simulations beforehand demonstrated, MAST is not necessarily a reliable “gold standard” to compare against. However, the 331 DE genes found by eSVD-DE are also much more enriched in GO terms that are plausibly related to ASD when compared to the DESeq2 genes or the original genes reported for layer 2/3 (Figure 5D). All these observations suggest that eSVD-DE is well-equipped to uncover meaningful results.

Next, we investigated qualitative differences among the methods compared to the SFARI and bulk DE genes. We enable fair comparisons among the different methods by finding the top 100 genes with the smallest p-values for each method. We then counted how many of these 100 genes intersected with the SFARI or bulk DE genes (Figure 5E). eSVD-DE has the highest overlap with both sets (31 with the bulk DE, 13 with the SFARI genes), followed closely by DESeq2 and then finally by MAST and SCTransform. In fact, eSVD-DE and DESeq2 have an overlap of 27 genes. The observations from Figure 5D and E combined suggest that eSVD-DE inherits many advantages of pseudobulk methods while improving these methods by accounting for within-individual variability. In the second investigation, we asked how stable each method was as the dataset was gradually downsampled. Indeed, as the scRNA-seq dataset becomes sparser, the relative difference between “strongly-expressed” DE genes and null genes becomes fainter, and we would expect any method to degrade in performance. To investigate this, we take the 50 genes with the largest test statistics in magnitude among the SFARI and bulk DE genes for each method and ask how their mean test statistic (relative to the housekeeping gene’s test statistic) diminishes as the scRNA-seq data is downsampled. We use the housekeeping genes as a proxy for the null genes, which is a reasonable choice given Figure 5B. As the downsampling percentage increases (i.e., the signal becomes fainter), MAST and SCTransform lose their ability to distinguish between formally highly-differentiable genes and house-keeping genes. This finding makes sense, as MAST and SCTransform estimate the DE genes based on a gene-by-gene regression, meaning no information is shared between genes. On the other hand, DESeq2 and eSVD-DE are surprisingly stable – they can demonstrate a clear separation between the 50 DE genes and the housekeeping genes even when the data is downsampled at 40%. This finding is also sensible, as DESeq2 aggregates cells among individuals, and eSVD-DE pools information between genes; both strategies yield robustness against sparsity. However, eSVD-DE is still preferred over DESeq2 here, as the separation between the DE genes and housekeeping genes is much higher for eSVD-DE.

We include additional diagnostic plots via Isomaps, volcano plots of the DE genes, and the GO analysis results in Appendix (Supplementary Figure F7, and F10 through F12).

## Conclusions

We demonstrate the nuances of performing cohort-wide differential testing for single-cell RNA-seq data. The difficulty primarily stems frotm individual-level confounding covariates, which can be difficult to remove using typical DE strategies of regressing out their effects gene-by-gene. Instead, eSVD-DE pools the information across genes to remove the confounding covariates, yielding empirical performance that is more promising than current pseudobulk DE methods or DE methods currently used for scRNA-seq data when no prevalent individual-level covariates are present. We achieve this through a matrix factorization strategy, estimating the coefficients associated with the covariates and each cell’s and gene’s latent vectors. We then shrink the estimated denoised gene expressions via the posterior mean according to the Gamma-Poisson distribution. This strategy helps dampen potential over-smoothing effects induced by the matrix factorization. We then deploy a test statistic designed to test for DE genes on the individual level instead of the cellular level. Note that our procedure can be used concurrently with IDEAS [27], which aims to find differential *distributions* between cases and controls, which could be more challenging to interpret than differential means. However, we do not pursue this direction in this paper. We also note that gene selection for cohort-level scRNA-seq datasets is an important task that we are interested in exploring in future work, since the inclusion of certain genes could impact the p-value of other genes due to the nature of pooling information across cells and genes via a low-dimensional embedding. Ideas from graph-representation work such as [51] and [52] could be highly relevant in this direction. We hope that eSVD-DE would be beneficial for inspiring cohort-wide DE tests for future single-cell assays beyond scRNA-seq and single-cell eQTL analyses where abundant individual-level covariate effects must be adequately removed. Additionally, we are curious about broader settings where instead of having case and control individuals within a cohort, we are interested in testing if continuous covariates such as age have a substantial transcriptomic impact on specific cell types, as discussed in [53].

### Statistical model and method

Let *A* ∈ { 0, 1, … }, ^*p×n*^ denote the observed count matrix with *n* cells and *p* genes, and *C* ∈ ℝ^*n×r*^ denote the observed *r* covariates for the *n* cells. Importantly, certain columns of *C* would be the following:

- **Intercept**: Let *C*_.,1_ = 1. We’ll call this column *C*_.,(int)_.
- **Log sequencing depth**: Let 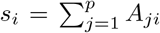. Then let *C*_*i*,2_ = log(*s*_*i*_) for all *i* ∈ {,…,*n*} We’ll call this column *C*_.,(lib)_.
- **Case-control status**: Let *C*_*i*,3_ ∈ {0, 1} depending on whether or not cell *i* ∈ {1, …, *n*} is associated with a case or control individual. We’ll call this column *C*_.,(cc)_.
- **Others**: The remaining columns of *C* could contain numerical covariates associated with the cells: which region of the body the cell was sampled from, the age of the corresponding individual, etc. We assume the categorical covariates have already been transformed via one-hot encodings (i.e., categorical variables with *k* levels are transformed using *k* − 1 indicator variables). In contrast, assume the numerical covariates are standardized to have a standard deviation of 1. We have found it beneficial to not center the numerical covariates around 0. This way, the resulting estimated coefficients in *Z* are easier to interpret.

Optionally, practitioners may include one-hot encoding vectors for which cells originate from which individuals. In our paper, we have found this to be optional and sometimes detrimental to the fit. This is because the inclusion of such one-hot encoding vectors result in a collinear matrix *C*. This preparation of the covariate matrix *C* is primarily handled by the function `eSVD2::.reparameterization esvd covariates` in our codebase.

The statistical foundation of eSVD-DE is the following hierarchical model where each entry of *A*_*ji*_ is modeled as,

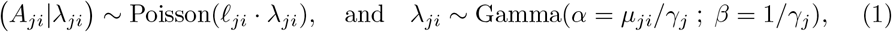

(where *α* and *β* denote the shape and rate parameters of a Gamma distribution respectively) for gene *j* ∈ {1, …, *p*} and cell *i* ∈ {1, …, *n*}, where the low-dimensional mean matrix is *µ* ∈ ℝ^*p×n*^ where

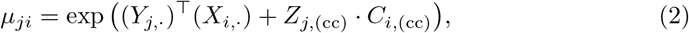

for *X* ∈ ℝ^*n×k*^ and *Y* ∈ ℝ^*p×k*^, and 𝓁_*ji*_ denotes the covariate-adjusted sequencing depth,

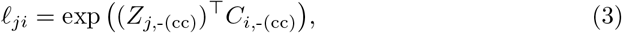

and *γ*_*j*_ *>* 0 denotes the overdispersion parameter for gene *j* which controls the variability of gene *j*. Here, the subset notation “(cc)” means that we exclude the “− (cc)” column from the corresponding matrix. In the terminology of previous work [22, 25], we interpret *µ*_*ji*_ as the “predictable” gene expression (i.e., expression of gene *j* in cell *i* that can be predicted from other cells and genes, which we interpret as a low-dimensional manifold), and *λ*_*ji*_ as the biological relative expression of gene *j* in cell *i*. We observe the count matrix *A* as well as the covariates *C*, and we need to estimate the cells’ latent embedding *X*, the genes’ latent embedding *Y*, the covariates’ coefficients *Z* as well as the overdispersion vector *γ*.

#### Regarding the Gamma distribution

Our parameterization of the Gamma distribution is inspired by the constant Fano factor model used in SAVER [22]. Specifically, for the Gamma distribution in (1),

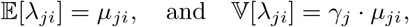

meaning the variance scales proportionally with the mean. Additionally, it can be derived that the marginal distribution of *A*_*ji*_ is a negative-binomial,

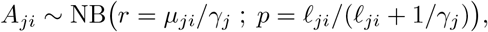

meaning the moments of *A*_*ji*_ marginally are

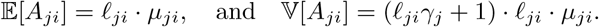

These equations demonstrate that *γ*_*j*_ measures overdispersion – the larger *γ*_*j*_ is, the larger the variance of *A*_*ji*_ is.

Additionally, we can derive that the posterior distribution of *λ*_*ji*_|*A*_*ji*_ is

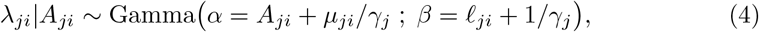

which will be useful when deriving our method below.

### Rationale and justification of statistical model

Our hierarchical model (1) draws inspiration from two literatures. The first literature is the single-cell modeling literature. Various work have shown that hierarchical model (1) that model observed counts as a Poisson distribution originating from a Gamma prior is a statistically sound and empirically justified model of scRNA-seq data [22, 25]. These models are appealing as they explicitly separate the technical noise (i.e., noise incurred by the nature of sequencing, modeled via the Poisson distribution) from the biological noise (i.e., noise naturally occurring among cells, modeled via the Gamma distribution). Other work has provided additional biological explanations on why biological noise is commonplace in organisms [54].

The second literature is the differential-expression literature, which canonically focuses on testing for the differential expression of a gene, one gene at a time. For gene *j*, methods like DESeq2 [8], MAST [10], and SCTransform [39] regress the observed expression *A*_*j*1_, …, *A*_*jn*_ onto observed sequencing depth of each cell 𝓁_1_, …, 𝓁_*n*_. These models account for the possibility that the sequencing depth detrimentally confounds the differential expression of gene *j* in varying degrees across all genes. Hence, our modeling of the sequencing depth in (3) is qualitatively similar.

### High-level description of the hypothesis test

We provide a high-level description of the hypothesis that eSVD-DE tests, with respect to the hierarchical model prescribed in the aforementioned statistical model. Let *A* denote the set of case individuals and *B* denote the set of control individuals. Consider a gene *j* ∈ {1, …, *p*}, assume that all there are parameters 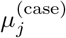 and 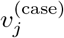 that depict the unknown gene expression mean and variance among the case individuals, i.e.,

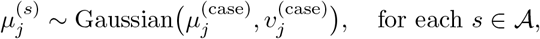

and likewise for the control individuals for unknown parameters *µ* ^(control)^and *σ*^(control)^,

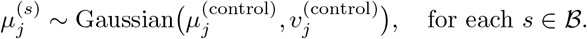

Each individual then contributes multiple cells. Let ℐ (*s*)⊂ {1, …, *n*} denote the set of cells that an individual contributes. We model cell *i*’s expression (from individual *s*, where *s* refers to “subject”) for gene *j* as

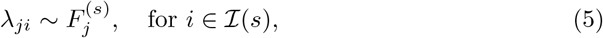

where 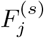 is a distribution with mean 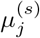. From here, the observed count in cell *I* for gene *j*, defined as *A*_*ji*_ is related to *λ*_*ji*_ through (1).

With this alternative perspective of our statistical model, eSVD-DE is testing between the null and alternative hypotheses,

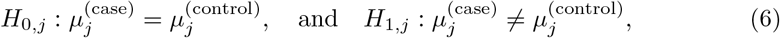

which are comparing the pouplation case-individual mean expression to the pouplation control-individual mean expression of gene *j*.

A few notes are in order:

- **Rationale of against pseudobulk approaches**: The hypothesis test in (6) might suggest a pseudobulk approach (such as used by DESeq2 in our comparisons). However, as we have mentioned in the main text, such pseudobulk methods do not capture variability within an individual (i.e., the variance of the distributions 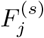. This can affect validity of such pseudobulk strategies, as demonstrated in our null simulations shown later.
- **Rationale of our hypothesis test**: We note that our hypothesis test in (6) relates the population case individual’s against the population control individual’s mean expression. This is different from the following null hypotheses that are not as apt for testing for differential mean expression for cohort-wide scRNA-seq data:

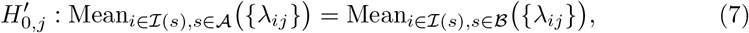

or

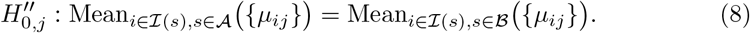

or We cannot test the null hypothesis in (7) directly since we observe only one observation *A*_*ji*_ for each *λ*_*ji*_. By pooling all the information across cells and genes, we can instead estimate *µ*_*ji*_, which is the mean of *λ*_*ji*_ based on a low-rank matrix factorization. In contrast, (8) ignores which cells originated from which individual, which makes it undesirable for differential testing for cohort-wide scRNA-seq data. For example, if all the cells from a few case individuals have vastly higher gene expression than all other case individuals’ cells, the null hypothesis in (8) might be rejected, but eSVD-DE’s null hypothesis in (6) would not be rejected. In such a scenario, we would not want to reject the null hypothesis since the behavior of a small number of case individuals would not be representative of the human population of case individuals. In our comparisons, methods like SCTransform test the null hypothesis in (8). Methods like SCTransform face a different computational obstacle. Since these methods regress out the individuals’ substantial covariate effects one gene at a time, this regression might be inaccurately estimated due to the sparsity of scRNA-seq data.
- **Distinction between cell-mean and individual-mean**: Observe that even though our hypothesis testing framework in (5) treats all the *λ*_*ji*_’s for the same individual (i.e., *i* ∈ *I*(*s*)) as i.i.d., eSVD-DE nonetheless models each cell’s mean as *µ*_*ji*_ in (2). This is to facilitate the matrix factorization framework in order to pool information across cells and genes. As we will discuss later in the model, we estimate 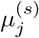 by averaging the estimated *µ*_*ji*_’s (after taking its posterior distribution to account for model misspecification, see (14)). We believe it is important to rely on the posterior distribution especially for cohort-wide differential expression testing since single-cell sequencing data is quite sparse, so it’s important to have a data-driven procedure to adjust our estimated values of *µ*_*ji*_ that balances how sparsely sequenced (or said differently, how much “information”) gene *j* is and how large the covariate-adjusted sequencing depth of cell *i* is.
- **Covariate effects, sparsity, and overdispersion**: Within this perspective of the model, all the confounding covariate effects is modeled through the covariate-adjusted sequencing depth 𝓁_*ji*_ in (3), which eSVD-DE tries to model appropriately by pooling information across cells and genes. Additionally, we lose power to test for (6) as sparsity increases, which is parameterized by the overdispersion parameter *γ*_*j*_.

### Implementation and details of eSVD-DE

#### Initialization

The initialization broadly falls into three steps: 1) initializing the estimate of the coefficients 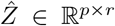, 2) initializing the estimate of the embeddings for the cells 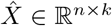 and for the genes *Ŷ* ℝ^*p×k*^, and 3) reparameterizing our estimates 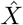, *Ŷ*and 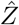. We describe each step below.

First, to initialize our estimate 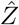, we fit a Poisson regression with a ridge penalty separately for each cell *i*. This regresses the covariates *C* onto *A*_.,*i*_, the observed gene counts for cell *i*. Specifically, for a small penalty *τ >* 0, we initialize the *j*th row of 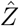 as

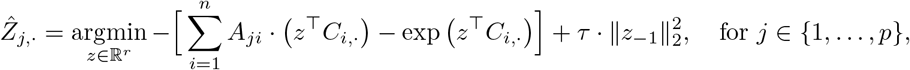

where *z*_−1_ = (*z*_2_, …, *z*_*r*_) ∈ ℝ^*r*−1^. The ridge penalty omits *z*_1_ because, by definition, *C*_.,1_ is the all-one vector that represents the intercept. Typically, we set *τ* = 0.01, a small non-negative value to mitigate multicollinearity issues, similar to other works such as [12].

Second, we initialize our estimates 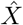 and *Ŷ*. Consider the matrix,

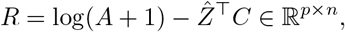

where log(·) of a matrix denotes taking the natural logarithm of each matrix entry. Consider the SVD of *R*,

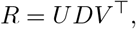

where *U* ∈ ℝ^*p×k*^ and *V* ∈ ℝ^*n×k*^ are column-wise orthonormal matrices, and *D* is a diagonal non-negative matrix with decreasing values along the diagonal. We then initialize our estimates of 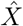 and *Ŷ* as

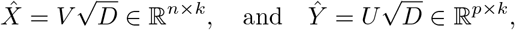

where 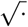 of a diagonal non-negative matrix denotes taking the square root of all its diagonal entries. This initialization of 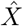 and *Ŷ* is motivated by the observation that if we model *A*_*ji*_ as a Poisson random variable, then log(*A*_*ji*_ + 1) is a crude approximation of the natural parameter *µ*_*ji*_, and 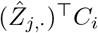_,*·*_would be an initial estimate of the covariate effect on *µ*_*ji*_.

This step is collectively handled by `eSVD2::initialize esvd` in our codebase. It relies on the `glmnet::glmnet` function to fit the Poisson ridge regression.

### Optimization of the embeddings

The iterative optimization of the embedding broadly falls into two steps to estimate the latent factors of the mean matrix *µ* defined in (2): 1) updating the estimates of 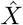, *Ŷ*, and 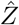, holding the initialized values of 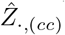 fixed, and then 2) updating the estimates of 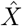, *Ŷ*, and 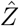, allowing the values in 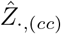 to be updated. This step takes inspiration from methods such as GLM-PCA [12], ZINB-WaVE [13], and our previous work [18]. We first describe the details of the procedure, followed by its justification.

For a tuning parameter *τ >* 0, we seek to optimize the following objective function

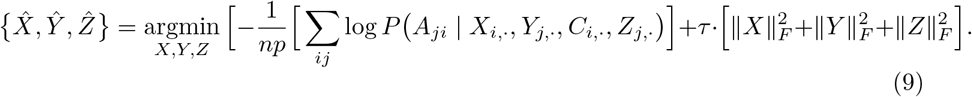

Let us define the natural parameter,

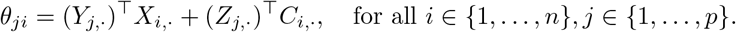

Then, here, log *P* (*A*_*ji*_ | *X*_*i*,._, *Y*_*j*,._, *C*_*i*,._, *Z*_*j*,._) denotes the log-likelihood for a Poisson random variable,

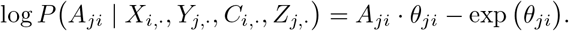

The objective function in (9) is a non-convex, as documented in [18]. This poses challenging optimization considerations. Hence, we first describe an alternating minimization strategy that improves upon previous work computationally. Then, we describe the aforementioned two-step approach.

Our alternating minimization consists of the following steps, starting on an initial estimation of 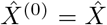, *Ŷ* ^(0)^ = *Ŷ*, and 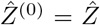. This strategy is motivated by the observation that holding *Y* and *Z* fixed, the optimization in (9) over *X* is convex, and vice-versa. Let *t* denote the iteration counter.

- Optimize the cell embedding,

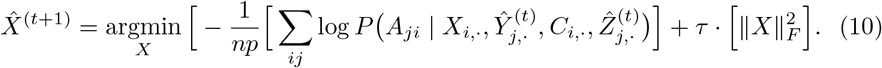
- Optimize the gene embedding and coefficients

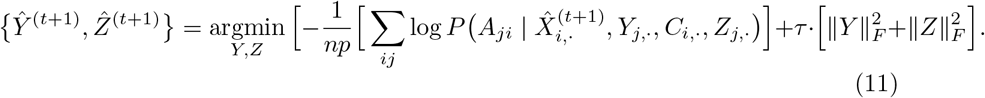

We repeat until convergence and then perform a reparameterization (described in the next section).

The optimizations (10) and (11) are performed using a Newton optimization, which is a second-order method. This specific optimization framework is ideal for solving (10) and (11) for a few reasons:

#### 1. Parallelization into many low-dimension optimizations

Both optimizations (10) and (11) decompose into *n* and *p* smaller optimization problems respectively.

For example, the *i*th row of 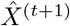 is equivalently solved by the optimization

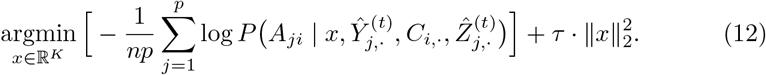

Hence, solving (10) and (11) amounts to a solving *n* different *K*-dimensional or *p* different (*K* + *r*)-dimensional optimizations respectively. Since we are in a setting where max(*K, r*) ≪ min(*n, p*), it is beneficial to use a second-order method since the Hessian information can yield faster convergence rates in terms of the number of iterations needed to solve optimizations like (12), and there is not a large computational overhead to compute the Hessians to solve (12) (especially when *P* is the Poisson distribution, where the Hessians are straight-forward to derive and compute).

#### 2. No randomness in the optimization

Since (10) and (11) decompose into *n* and *p* smaller optimization problems respectively, these optimizations can be embarrassingly parallelized. Hence, there is no need to consider stochastic optimization schemes. This is appealing as this means different practitioners using our method would necessarily obtain the same resulting fit.

#### 3. Generalization to other exponential families

While we focus on specifically solving (10) and (11) for the Poisson distribution, using a Newton optimization is ideal for other exponential-family distributions as well. This is because while other exponential-family distributions like the exponential and Negative Binomial have constraints on the natural parameters *θ*_*ij*_, the gradient and Hessians of these log-likelihoods naturally prevent the optimization iterates from violating these constraints. Hence, our codebase can handle modeling situations beyond this paper’s scope.

Our optimization procedure is then the following. First, starting from the current estimates of 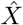, *Ŷ*, and 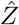 in the previous step (“Initialization”). we updating the estimates of 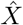, *Ŷ*, and 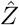 via alternating minimization holding the initialized values of 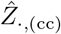 fixed (what we’ll call the “Phase one optimization”). After convergence, we then perform a second round of alternating minimization to update the estimates of 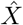, *Ŷ*, and 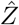, allowing the values in 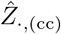 to be updated (what we’ll call the “Phase two optimization”). (Here, we omit the superscript “(*t*)” for notational simplicity.) We do these two phases of optimization since empirically, we have observed more stable behavior and a better final objective value doing two rounds of optimization (compared to optimizing all of *X, Y*, and *Z*, including *Z*_.,(cc)_ from the start). This becomes imperative since *Z*_.,(cc)_ will play a more pronounced role in the remainder of eSVD-DE compared to all the other covariates in *Z*, because is it part of the “signal” that we wish to estimate and is not a “confounder.” This is inspired by theoretical results regarding warm-starting the non-convex optimization [55, 56]. This step is collectively handled by `eSVD2::opt esvd` in our codebase.

### Reparameterizing the matrix factorization

After completing either of the two phases mentioned above, we apply the following reparameterization procedure. The goal of this reparameterization is to ensure identifiability since, a priori, there could be multiple estimates 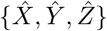 that have the same predictive power but offer different interpretations of the data. For instance, without such a reparameterization, there could be many columns in 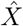 that are correlated with other columns in 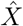 or columns in the covariate matrix *C*, which would obfuscate interpreting different axes of variation. Hence, our reparameterization procedure ensures orthogonality among our estimated matrix factorization to resolve this identifiability concern.

When describing our reparameterization procedure, for notational simplicity, we let { 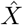, *Ŷ*,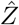} denote {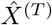, *Ŷ* ^(*T*)^,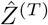}, which is the final estimate after *T* iterations for either the Phase one or two optimizations. We also let the “*a* ←*b*” notation denote setting the variable *a* to be the value in variable *b*.

The reparameterization procedure operates in two steps. The first step ensures that 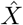 is orthogonal to *C* (which would result in adjusting 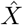 and 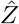). This ensures that 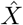 captures variability that is not explained by *C*. The second step ensures that both 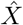 and *Ŷ* have orthogonal columns (which would result in adjusting 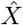 and *Ŷ*). This ensures that each respective latent dimension in both 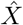 and *Ŷ* are identifiable.

- **Step 1**: We first perform a regression of each latent dimension of 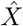 onto *C*. That is, for each latent dimension *d* ∈ {1, …, *k*},

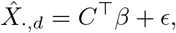

where *β* ∈ ℝ^*r*^ and *ϵ* ∈ ℝ^*n*^ are the temporary variables to denote the coefficients for the covariates and residuals respectively. For this latent dimension *d*, we then perform the following update for each gene *j* ∈ {1, …, *p*},

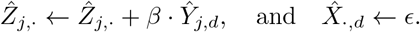

It can be seen that after performing this update for every latent dimension 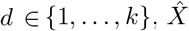 is orthogonal to *C* (i.e., 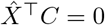) even though the predictive power of our factorization (i.e.,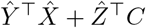) did not change.
- **Step 2**: We next perform a linear transformation on 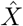 and *Ŷ*. Specifically, using the details in [18], let 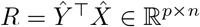, with an SVD of *R* = *UDV* ^⊤^. Then,

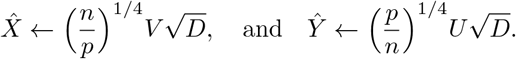

It can be seen that after performing this update that both 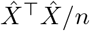 and *Ŷ*^⊤^ *Ŷ/p* are diagonal matrices and are equal even though the predictive power of our factorization (i.e., 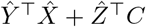) did not change. This ensures identifability of 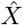 and *Ŷ*.

### Estimating overdispersion parameter

We estimate the overdispersion parameter *γ*_1_, …, *γ*_*p*_ *>* 0, one for each gene. Following our model of the covariate-adjusted sequencing depth (3), we estimate the covariate-adjusted sequencing depth of each cell as

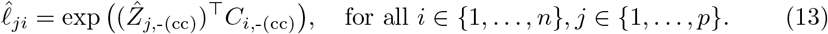

Likewise, the estimate of the mean parameter is

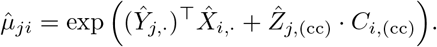

Then, using a plug-in estimate of the maximum likelihood of the model in (1), we can derive that the estimate of *γ*_*j*_ is

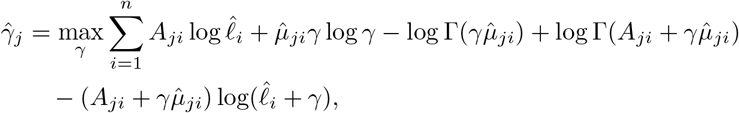

where Γ(.) is the Gamma function. We estimate this via Newton’s method, and this step is primarily handled by `eSVD2::estimate nuisance` in our codebase.

### Computing the posterior distribution

To account for possible model misspecification, we estimate the posterior distribution of the mean and variance of each gene *j*’s expression in cell *i*. This is based on the model in (1), where we can leverage the fact that *λ*_*ji*_ conditioned on *A*_*ji*_ follows a Negative Binomial distribution. Specifically, based on the posterior distribution derived in (4), we compute

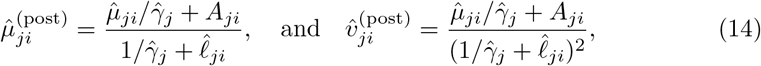

where 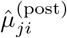 and 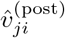 is the posterior mean and variance of *λ*_*ji*_|*A*_*ji*_ for gene *j* in cell *i* respectively. Other work have used similar modeling based on the posterior distribution for single-cell RNA-seq data [22], but not for the purposes of DE testing. This step is primarily handled by `eSVD2::compute posterior` in our codebase.

### Computing the test statistic

In this step, we compute the test statistic, accounting that multiple cells originate from a particular individual. This broadly falls into two steps: 1) computing the mean and variance for a gene among an individual, and 2) computing the test statistic among all the case and control individuals.

With the posterior distribution in (14), we can compute the expected posterior mean and variance for a particular gene *j* in individual *s*. First, we aggregate among the cells for each individual. We assume that after averaging among all the cells from an individual, the resulting distribution can be reasonably approximated by a Gaussian with mean 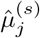 and variance 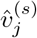. Hence, let *ℐ* (*s*) ⊂ {1, …, *n*} denote the set of cells originating from individual *s*. Then, via large-sample average,

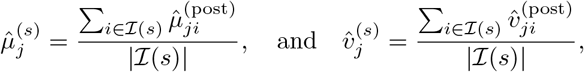

where 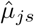 and 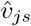 is the posterior mean and variance for gene *j* among all the cells in individual *s* respectively.

Next, we aggregate among individuals. Specifically, among all the case individuals, we can think of the expression of gene *j* as a Gaussian mixture among the individuals. Let 𝒜 denote the set of case individuals. We then summarize the Gaussian mixture with one Gaussian,

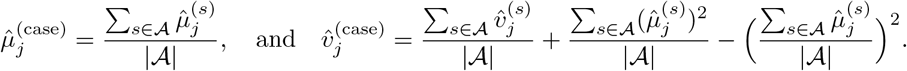

We do an analogous calculation among controls, letting ℬ denote all the control individuals.

Lastly, we do a two-sample T-test with unequal variances, treating the distribution among cases as a Gaussian with mean 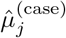 and variance 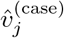 and the distribution among controls as an analogous Gaussian. Our test statistic for gene *j* is

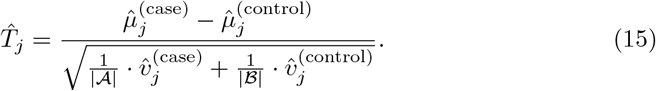

In our codebase, this step is primarily handled by `eSVD2::compute test statistic`.

### Performing multiple testing correction

We use the multiple testing procedure based on the empirical null developed in [57] to handle the multiple testing corrections. This framework is also used in other methods designed to test for differential expression from scRNA-seq data (albeit not from cohort data), such as iDEA [58] and SifiNet [59]. We know that biologically, most genes are not directly related to the disease or disorder at the human population level; hence, most genes should be deemed insignificant. The empirical null, where the appropriate null distribution is learned from the data, is a suitable framework to ensure this behavior.

Briefly, we first estimate the degree-of-freedom of each gene *j* ∈ {1, …, *p*} (used for T-tests for unequal variances) by

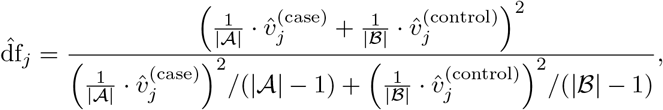

and convert all the test statistics 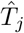’s into a Z-score via,

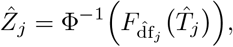

where *F*_*b*_(*a*) is the CDF of the a *t*-distribution with degree freedom *b >* 0 evaluated at *a*, and Φ^−1^(*a*) is the quantile function of a standard Gaussian evaluated at *a* ∈ (0, 1).

To identify the DE genes, we use the `locfdr::locfdr` function to estimate the empirical null distribution’s mean and standard deviation. We then compute the p-value of each gene based on the (two-sided) tail area of a Gaussian distribution with that empirical null’s mean and standard deviation. We can compute the negative log_10_ p-value for visualization purposes in a volcano plot (as in Figures 3 and 4). To perform multiple testing corrections, we apply `stats::p.adjust` on these p-values via Benjamini-Hochberg. All the genes with an empirical FDR of less than 0.05 are then deemed to be DE genes.

This step is collectively handled by `eSVD2::compute df, eSVD2::compute test statistic`, and `eSVD2::compute pvalue` in our codebase.

### Information about data preprocessing

We describe the high-level ideas on what is needed for our preprocessing of the data prior to using the eSVD-DE. Importantly, we need to: 1) select the cells specific for our analysis, and 2) select the genes of interest for our analysis. For all the datasets in our analysis, the cell type labels were already provided by the author, and we used `Seurat::FindVariableFeatures` to select most of the genes in our analysis. (We also included housekeeping genes and genes previously reported to be differentially expressed by the authors as well.) No preprocessing of the count data is needed as eSVD-DE models the raw count data directly, using the observed sequencing depth and covariates of the individuals. We more thoroughly detail these steps in the Appendix (Section E) and include details on which covariates were included in all our analyses as well as the parameters used for eSVD-DE.

#### Supplementary information

High-level description on what is statistically challenging about doing differential expression testing after denoising a datset, simulation details and results, and supplemental details of the lung, colon, and brain analyses, and supplemental figures A1-F12.

Additionally, `adams-habermann-T esvd results.xlsx` contains the analysis of T-cells of either the Adams data (from [6]) or the Habermann data (from [19]). It contains the estimated log-fold change after removing confounding effects, the eSVD-DE test statistic, the negative log_10_ p-value, and Benjamini-Hochberg adjusted p-value, and whether the gene is deemed to be a DE gene. Similarly, `smillie esvd results.xlsx` contains the same 5 columns, but for cycling TA, enterocyte progenitors, TA 1, and TA 2 when analyzing either inflamed or non-inflamed tissues in the Smillie dataset (from [20]). Lastly, `velmeshev esvd results.xlsx` contains the same 5 columns, but for astrocytes, endothelial cells, IN-SST, IN-VIP, layer 2/3, layer 4, layer 5/6, layer 5/6-CC, microglia, oligodendrocytes, and OPC when analyzing the Velmeshev dataset (from [21]).

## Acknowledgments

We thank Timothy Barry, Jing Lei, Ronald Yurko, and Nancy Zhang for useful comments when developing this method.

## Declarations

### Funding

This project was funded by National Institute of Mental Health (NIMH) grant R01MH123184.

### Competing interests

The authors declare that they have no competing interests.

### Ethics approval

Not applicable

### Consent to participate

Not applicable

### Consent for publication

Not applicable

### Availability of data and materials

All analyses were done on publicly available data. The Adams data (from [6]) is downloaded from GSE136831, and the individual covariates were provided by the authors of the paper. The Habermann data (from [19]) is downloaded from GSE135893, and the individual covariates were provided by the authors of the paper. The Smilie data (from [20]) is downloaded from https://github.com/cssmillie/ulcerative colitis, while the individuals’ covariates and DE genes are downloaded as Table S1 and S4 from https://www.ncbi.nlm.nih.gov/pmc/articles/PMC6662628/. The Velmeshev data (from [21]) is downloaded from https://cells.ucsc.edu/?ds=autism, while the DE genes is downloaded as Data S4 from http://science.sciencemag.org/content/suppl/2019/05/15/364.6441.685.DC1. The housekeeping genes (from [40]) is downloaded as Supplementary Table 1 from https://academic.oup.com/nar/article/49/D1/D947/5871367#supplementary-data. The SFARI genes were downloaded from https://gene.sfari.org/database/gene-scoring/ on January 6, 2022 (using the September 2, 2021 release). The Gandal genes (from [49]) is downloaded as Supplementary Data 3 from https://www.nature.com/articles/s41586-022-05377-7, where we look at the DEGene Statistics sheet and select the genes with WholeCortex ASD FDR less than 0.05.

### Code availability

See https://github.com/linnykos/eSVD2 for the R package eSVD2 that contains all the functions used in for the eSVD-DE analysis. See https://github.com/linnykos/eSVD2 examples for all the scripts used in the analysis.

### Authors’ contributions

K.Z.L. and K.R. conceived of the idea. K.Z.L.. and Y.Q. coded eSVD-DE in R and C++. K.Z.L. performed the analyses. K.Z.L., Y.Q. and K.R. wrote the paper.

## Notes

### Competing Interest Statement

The authors have declared no competing interest.

### Summary of Updates

There were revisions primarily in the supplemental materials to bolster the robustness/power of the method shown through simulations. This is the current (Supplementary) Section C.2, C.3, D. Also added Section E.2 for data table summarizing the datasets in the paper.

