## Supplementary Materials for "eSVD-DE: Cohort-wide differential expression in single-cell RNA-seq data using exponential-family embeddings"

#### Appendix A Illustration of how dimension reduction introduces downstream Type-1 inflation in differential expression testing

Here, we provide a simplistic example of how dimension reduction complicates the validity of differential expression testing downstream. Our example relies solely on two genes across six individuals (three cases, three controls), where each gene's expression is Gaussian distributed. To focus on how dimension reduction interacts with downstream testing, we do not involve the count-nature of scRNA-seq data, sparsity of gene expression, or confounding covariate effects in this example.

The simple simulated data is shown in Figure A1A, where Gene 1 is differentially expressed whereas Gene 2 is not (two-sample test p-value of 0.0019 and 0.4723 respectively). Coincidentally, the direction of most variance is also strongly aligned with the axis of Gene 1. Hence, if we project the data onto the leading principal component, our resulting "denoised" dataset is shown in Figure A1B. When the same two-sample test is applied to each gene individually, both genes are differentially expressed (p-value of 0.0019 for both genes). Both genes have the same p-value in this example after projecting onto the first principal component since the "information" in both genes is identical (i.e., perfectly collinear).

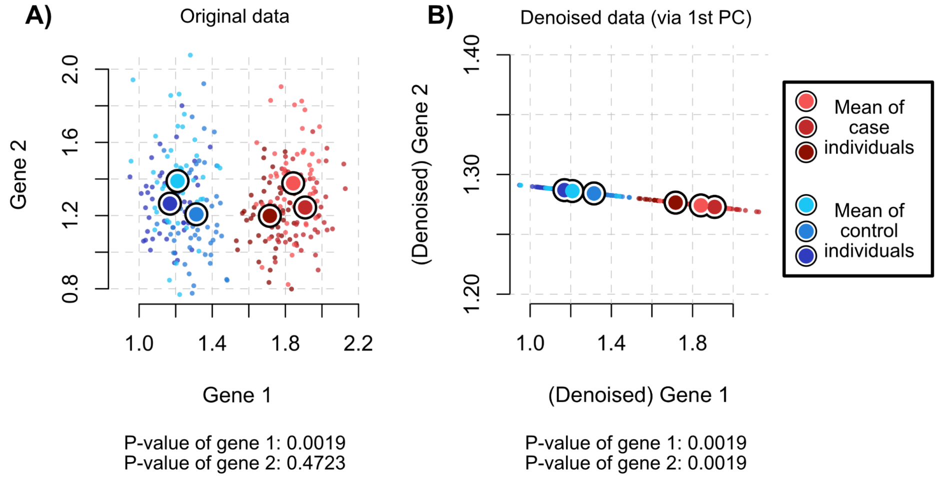


**Fig. A1:** **A** Scatter plot of the simulated dataset with multiple cells originating from six individuals (three cases and three controls) for two genes, where Gene 1 is designed to be a true DE gene (two-sample test p-value of 0.0019) and Gene 2 is not (p-value of 0.4723). **B** Scatter plot of the same dataset after all the cells are projected onto the leading principal component. Now, both genes appear to be statistically significant DE genes (p-values of 0.0019), illustrating how dimension reduction can inflate the downstream Type-1 error.

This example illustrates the following phenomenon. A dimension reduction (such as projecting data onto its leading principal components) will necessarily introduce correlation among all genes. When true DE genes primarily drive the matrix factorization, many true null genes will appear as statistically significant DE genes downstream after the dimension reduction. In many biological datasets, this is likely the scenario since a few DE genes often have a high signal-to-noise ratio, and the practitioner is interested in finding other DE genes with a weaker signal-to-noise ratio. However, after applying a dimension reduction, most genes will appear to have a DE gene with a weak signal-to-noise ratio.

In our particular example, it was clear that such a dimension reduction was not necessary for our goal of testing for DE genes. However, when there are thousands of genes (instead of two) that are sparsely observed count data (instead of densely observed Gaussian data) that are confounded by various covariates (instead of not having any covariate effects), dimension reduction is likely necessary to pool information across cells and genes. In that setting, the phenomenon we illustrate in Figure A1 is likely to happen still but would not be as easily detectable. This illustration highlights the necessity to consider the posterior distribution, which we describe later in (14) as a way to account for model misspecification by balancing how sparsely sequenced gene $j$ is with how large the covariate-adjusted sequencing depth of cell $i$ is.

We note that work using deep variational autoencoders like [1, 2] have developed another way to circumvent this problem. First, the nature of the variational autoencoder relies on simulating Gaussian data after encoding the single-cell gene expression profile (hence, "variational"). This necessarily weakens the correlation among genes despite reducing the dimensionality, in contrast to a matrix-factorization approach like principal component analysis. Second, they advocate adding a "pseudcount" to all the gene expressions, which filters out genes that are deemed as significantly differentially expressed but have a near-zero log-fold change. While both qualities provide a sound solution to perform DE testing after a dimension reduction, this strategy relies on a "black-box" method with numerous tuning parameters to train the deep neural network and to perform the DE test. We instead choose to develop a strategy reliant on the posterior distribution in eSVD-DE as a more transparent and tuning-free strategy.

### Appendix B Null simulation

We perform the following simulation to demonstrate that eSVD-DE properly controls the Type-1 error. To effectively illustrate this, we construct a simulation where out of the $p = 1000$ simulated genes, 10 of which are truly differentially expressed. The remaining 990 genes are null, and we are interested if the p-values for these 990 genes are uniformly distributed in 0 and 1. We then contrast our method's results with other methods commonly used to analyze cohort-wide single-cell data.

***Simulation setup.***

We simulate data among $p = 1000$ genes across 10 case individuals and 10 control individuals where each individual contributes 100 cells, forming a combined $n = 2000$ cells. The individual covariates we consider are sex (categorical, 5 males and 5 females in each case and control) and age (numerical, drawn from a Gaussian distribution with a mean of 30 and standard deviation of 5). These are encoded to the covariate matrix $C \in \mathbb{R}^{n\times 5}$, where the 5 covariates are the intercept (i.e., a column of all 1's), the log sequencing depth (where it temporarily set to all 0's), and the case-control status, the sex, and age of the cell's individual.

We first describe the generation of the true natural parameter matrix. For the 1000 genes, as mentioned before, the first 10 genes are truly differentially expressed. We simulate the natural parameter matrix $\Theta^* \in \mathbb{R}^{p\times n}$ gene-by-gene, where each gene falls in one of three categories:

1. **True DE gene**: For genes $j \in \{1, 2, \ldots, 10\}$, the mean value among all the case individuals is uniformly chosen between 0.5 and 3. The mean value among all the control individuals is set to be exactly 1 larger or smaller (randomly chosen) than the mean value among all the case individuals. For a group (i.e., either the cases or controls), the specific individual's mean value is sampled to be normally distributed around their respective group's mean with a standard deviation of 0.1. Then, for an individual, all cells originating within an individual are sampled to be normally distributed around the individual's mean with a standard deviation of 0.1.

   The result is a substantial difference between all the case and control individuals' values for this gene $j$. Figure B2(A,D) shows an example of such a gene.

2. **Null gene with imbalanced variance**: For genes $j \in \{11, 13, \ldots, 999\}$, the mean value among all individuals is uniformly chosen between $-0.5$ and 1. For both cases and control individuals, the specific individual's mean value is sampled to be normally distributed around their respective group's mean with a standard deviation of 0.1. Then, one group (i.e., the case or control, randomly chosen) is selected so all the individuals in that group have a within-individual standard deviation set to be 0.75 (i.e., high within-individual variation). In contrast, the other group has all the individuals have a within-individual standard deviation set to be 0.1 (i.e., low within-individual variation). Then, for an individual, all cells originating within an individual are sampled to be normally distributed around the individual's mean with their prescribed standard deviation.

   The result is that even though there is no difference in mean between all the cases from all the controls in aggregate for this gene $j$, the within-individual variation in one group is substantially higher than the other group. Figure B2(B,E) shows an example of such a gene.
3. **Null gene interleaved between case and control**: For genes $j \in \{12, 14, \ldots, 1000\}$, the mean value among all individuals is uniformly chosen between 0.5 and 2. For all individuals, the specific individual's mean value is sampled to be normally distributed around their respective group's mean with a standard deviation randomly chosen between 0.1 and 0.2. Then, for an individual, all cells originating within an individual are sampled to be normally distributed around the individual's mean with a standard deviation randomly chosen between 0.1 and 0.2.

   The result is that there is no difference between case and control individuals for this gene $j$. Figure B2(C,F) shows an example of such a gene.

Then, we threshold the entries in $\Theta^*$ to not be larger than $\log(100)$, then uniformly shift all the entries down to encourage more sparsity in the generated observed matrix downstream.

We then describe how to introduce the sequencing-depth effects based on the covariates. Due to how the covariate matrix $C$ is constructed, the only relevant columns of the coefficient matrix $Z^* \in \mathbb{R}^{p \times 5}$ that need to be discussed are the coefficients for the intercept (which are set to all 0), the sex (which are sampled to be normally distributed with mean 0 and standard deviation 0.2), and the age (which are sampled to be normally distributed with mean 0 and standard deviation 0.5). Then, we construct

$$\ell^*_{ji} = \exp\big((Z^*_{j,\cdot})^\top C_{i,\cdot}\big).$$

We then describe the generation of the overdispersion parameters. The overdispersion parameter $\gamma^*_j$ is set to be 1 for all truly DE genes (i.e., $j \in \{1, \ldots, 10\}$), and then uniformly randomly sampled among 0.1, 1, and 10 for all remaining 990 null genes.

Lastly, we describe the generation of the observed simulated data. Equipped with all the above ingredients, we simulated data according to the hierarchical model in (1). Specifically, letting the mean value of gene $j$ in cell $i$ be

$$\mu^*_{ji} = \exp\big(\Theta^*_{ji}\big),$$

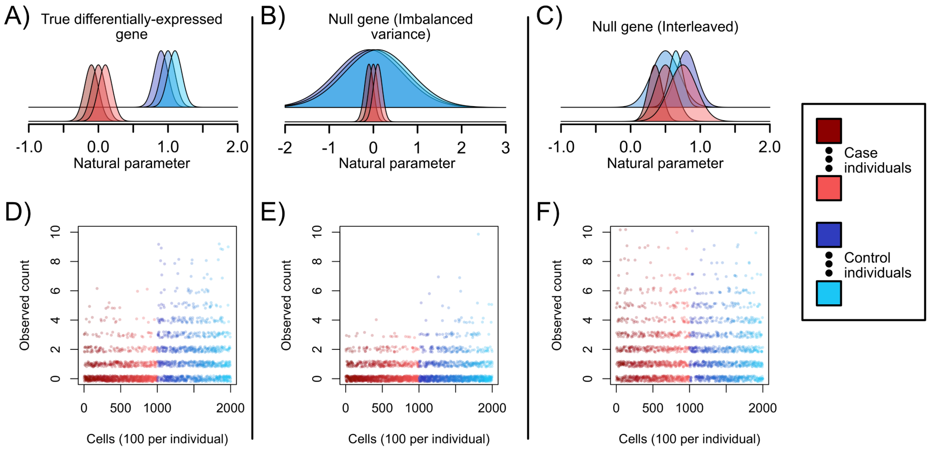


**Fig. B2**: **A** Density of values for a true DE gene, where there is a significant difference in means between the population case individuals and population control individuals, even when accounting for the within-individual variance. **B** Density of values for a null gene where there is a much higher within-individual variance among the control individuals than the control individuals, but the population mean among the case individuals' means is equal to the population mean among the case individuals' means (i.e., null gene with imbalanced variance). **C** Density of values for a null gene where the distribution of any individual's cells' expression is drawn from the same distribution (i.e., interleaved null gene). **D** Observed counts for a true DE gene across the six individuals when there are no confounding covariate effects, displaying a clear higher expression among the control individuals than the case individuals. **E** Observed counts for a null gene with imbalanced variance across the six individuals when there are no confounding covariate effects, displaying a larger variance among the control individuals than the case individuals. **F** Observed counts for an interleaved null gene, where both case and control genes have an unobservable difference in mean or variance. In all six figures, the simulated data displays the gene expression from three case individuals (in different shades of red) and three control individuals (in different shades of blue).

we sample the observed data matrix $A \in \mathbb{Z}_{+}^{p \times n}$ to be,

$$\big(A_{ji} | \lambda^*_{ji}\big) \sim \text{Poisson}(\ell^*_{ji} \cdot \lambda^*_{ji}), \quad \text{and} \quad \lambda^*_{ji} \sim \text{Gamma}(\alpha = \mu^*_{ji}/\gamma^*_j \ ; \ \beta = 1/\gamma^*_j),$$

The resulting dataset is roughly 60% sparse (i.e., only 40% of the entries are non-zero). We then appropriately set the values in $C_{\cdot,(\text{lib})}$ to be the log sequencing depth of the observed data.

We rationalize our design of the simulated data for the following reasons:

- **Clear definition of DE gene**: By the way we generated the values in $\Theta^*$, it is clear that in population, only genes $j \in \{1, \ldots, 10\}$ are differentially expressed because all the other genes have the same group-wise mean between cases and controls.
- **Demonstration of scenario where pseudobulk is not ideal**: Due to the null genes with imbalanced variances (i.e., genes $j \in \{11, 13, \ldots, 999\}$), the higher variation within-individual in one group versus the others means that if only pseudobulk samples were considered, the analysis loses information on how variable the cells are within an individual. Since the observed values in $A$ are only non-negative integers, a higher variance within an individual typically results in a higher observed mean. This means in this analysis, the pseudobulk samples could mistake cells with a higher within-individual variance to have a higher within-individual mean. In contrast, an

analysis on the single-cell level like eSVD-DE can retain the information that the variation within the individual is large for downstream test statistic calculations.

- **Demonstration of the necessity of pooling information in the presence of sparsity**: Lastly, the sparsity of each gene means that regressing out the covariate effect gene-by-gene is likely to result in a suboptimal removal of covariate effects.

***Methods used.***

We compare our eSVD-DE method against four other methods described previously in the main text. These are: 1) DESeq2, as a prototypical method that illustrates analyzing the data via pseudobulk samples, 2) MAST, as a prototypical method that illustrates analyzing the data via mixed-effect models, 3) SCTransform followed by Wilcoxon test, as a prototypical method that illustrates analyzing the data when ignoring which cells originate from which individuals, and 4) GLM-PCA followed by the Wilcoxon rank sum test, which prototypical method that illustrates analyzing the data using existing dimension-reduction pipelines. Notably, we are *not* critiquing any particular method specifically, but instead the overarching strategy of either regression out the covariate effects one gene at a time or using a dimension-reduction method followed by a Wilcoxon rank sum test respectively.

The goal is 1) to ensure that the 10 true DE genes are indeed assessed to be differentially expressed, but more importantly, 2) to verify that the remaining 990 null genes have a uniform distribution of p-values.

***Results.***

When applying all 5 methods to the data, all 5 methods consistently rejected the null hypothesis for the 10 true DE genes (across multiple trials). This ensured that each method had sufficient detection power.

However, when analyzing the p-values for the remaining 990 null genes, each method varied in the amount of Type-1 error. The QQ plots of the null genes' p-values are shown in Figure B3 (i.e., both imbalanced variance and interleaved). We see that eSVD-DE correctly has no Type-1 inflation among the null genes due to its pooling of information across cells and genes in the matrix factorization framework but also the posterior correction to avoid model misspecification. DESeq2 and MAST were the next two methods with the least Type-1 error, which reflects why pseudobulk or mixed-effect models are often deployed to analyze cohort-wide scRNA-seq data currently [3–5]. DESeq2's Type-1 inflation occurred specifically the genes with imbalanced variance. This is expected, as pseudobulk methods do not capture the within-individual variability. While MAST's Type-1 error stemmed from its inaccuracy to regress out the covariate effect (due to its regression gene-by-gene), it performed better than SCTransform and GLM-PCA due to its mixed-effect nature. Lastly, SCTransform and GLM-PCA inflated Type-1 errors dramatically. Importantly, we are not critiquing these methods specifically but rather the pitfalls of using these overarching strategies when testing for DE in cohort-wide scRNA-seq data. The Type-1 inflation is due to its inaccuracy to regress out the covariate effect and/or testing for DE on the cell-level instead of at the individual-level, as discussed throughout the main text and appendix.

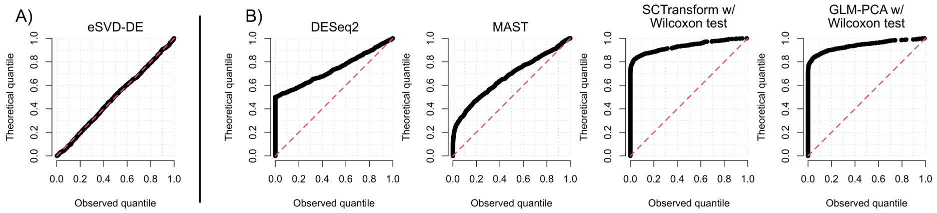


**Fig. B3**: **A** QQ-plot of the quantiles of the p-values for the null genes (theoretical vs. observed) for eSVD-DE on the null simulation in one trial. **B** For the same trial, the QQ-plots of the p-values for DESeq2, MAST, SCTransform (followed by a Wilcoxon rank sum test) and GLM-PCA (followed by a Wilcoxon rank sum test). In every plot, the red dotted line denotes the $x = y$ line. A method with no Type-1 inflation would show that the theoretical equal the observed quantiles among the null genes, which is only achieved by eSVD-DE in our null simulation.

### Appendix C Power simulation

#### C.1 Details of power simulation shown in main text

We now describe how the simulated data in Figure 2 was generated. The complexity of our data generation scheme stems from the desire to simultaneously estimate a gene's "true" log-fold change between cases and controls (excluding confounding effects) while also having genes of varying log-fold changes. We specifically do not want to simulate genes with explicitly no "true" log-fold change in this simulation. Instead, we wish to see if eSVD-DE's estimated log-fold change of a gene is strongly correlated with the "true" log-fold change of a gene. We also wish to make the data generation process complex so that it is misspecified with respect to eSVD-DE's assumed model. This quality helps to ensure that no method, not even eSVD-DE, has an unfair advantage when benchmarking eSVD-DE against other methods.

Since the data generation method is quite involved, please see the codebase for details. Below, we summarize the main aspects of the procedure.

Recall the model in (1). The first phase of data generation is to generate the cell embedding $X \in \mathbb{R}^{n\times k}$, the gene embedding $Y \in \mathbb{R}^{p\times k}$, the cell's covariate matrix $C \in \mathbb{R}^{n\times r}$, the coefficient matrix $Z \in \mathbb{R}^{p\times r}$, and the overdispersion vector $\gamma \in \mathbb{R}^p_+$. To do this, we simulate a population of 20 individuals (10 cases and 10 controls), each contributing 250 cells (meaning $n = 5000$ cells). The cell embedding $X$ is generated from $k$-dimensional multivariate Gaussians with non-trivial correlation, where $k = 10$. On the other hand, the gene embedding $Y$ by assigning genes to a weighted combination of 4 "topics" or no topics at all. For the genes assigned to the former category, the latent embedding for the gene is drawn from a simplex with 4 points (in a $k$-dimensional space). The latent embedding is drawn from a multivariate Gaussian centered at 0 for genes in the latter category. In total, there are $p = 700$ genes, where 200 of the genes are not assigned to any topic. For the covariate matrix $C$, the case-control status is set accordingly so the first 10 individuals are cases, and the last 10 individuals are controls. There is also a column for the intercept, a column for the log sequencing depth (which is set to be 0 for now), as well as the covariates for age (normally distributed), sex (boolean, equally distributed between cases and controls), and tobacco usage (boolean, correlated with the case-control status). Hence, $r = 6$.

The coefficient matrix $Z$ is constructed column-by-column, where the coefficient for a gene is set according to which topic the gene was assigned to, and different topics have different distributions of the coefficient value. Importantly, the coefficient for each gene for the case-control status of each cell is drawn among five different bins: "strong-negative," "weak-negative," "none," "weak-positive," and "strong-positive," for a coefficient drawn from a distribution with mean that is $-0.75$, $-0.5$, 0, 0.5 and 0.75 respectively. The coefficient for the log sequencing depth is set to be all 1's. Lastly, the nuisance parameter, taking values from 0.1, 1, and 100, is also set according to the topic the gene was assigned to, where, in general, if a gene has a larger case-control coefficient, we set the gene to be less overdispersed.

Equipped with all the components of the model, we can generate the low-dimensional mean matrix $\mu \in \mathbb{R}^{p \times n}$ according to (2). However, to make the model misspecified with respect to the eSVD-DE, we perform two additional modifications to the generated low-dimensional mean matrix $\mu$. First, depending on which genes had a large case-control coefficient in magnitude, we slightly shift all the 250 cells for each of the 20 individuals by a constant amount. This way, there is more variation among these genes that is not solely captured by the boolean case-control status vector in $C$. Second, for many of the genes with a small case-control coefficient, we artificially shrink the expression for that gene to become more correlated with the genes with a large case-control coefficient. This way, there is more correlation among the $p$ genes. We then generate the matrix $\lambda \in \mathbb{R}^{p \times n}$ and the observed count matrix $A \in \{0, 1, \ldots, \}^{p \times n}$ from the modified mean matrix $\mu$ and the overdispersion vector $\gamma$, following (1).

Once the observed count matrix has been generated, we can compute the observed log sequencing depth of each cell and set the column accordingly to $C_{\cdot,(\text{lib})}$. To assess the "true" gene expression not impacted by the confounding covariates, we regress $\mu$ onto all the columns of $C$ aside from the column for the case-control status, $C_{\cdot,(\text{cc})}$, and inspect the residuals. For these residuals, we can compute the "true" test statistic of each gene unaffected by the confounding covariates using calculations akin to those in the "Computing the test statistic" and "Performing multiple testing correction" subsections above. This lets us know the true log-fold change between case and control individuals for each gene. Also, we can compute the "true" FDR of each gene, and we deem all genes with a "true" FDR of less than 0.05 as a "true" DE gene.

#### C.2 Extension of power simulations

In order to determine how our method performs in a range of situations more realistic of ones observed with typical cohort-level scRNA-seq datasets, we vary the simulation along two axes – changing the number of genes, and changing how unevenly distributed the cells are across the 20 individuals. We consider three different number of genes – 100, 700, and 5000. By changing the number of genes, we are able to investigate how many genes are needed for eSVD-DE's low-dimensional embedding to be beneficial. (If there are not many genes in the analysis, one can imagine that eSVD-DE's low-dimensional embedding offers little practical advantage over other methods.) We consider three different amount of "unequalness" in the number of cells for each individual – equal distribution of cells in each individual, a maximum difference of

4x number of cells between any two individuals, and a maximum difference of 16x number of cells between any two individuals. Specifically, the last setting means there exists two individuals where one individual has 16 times more cells in the dataset compared to another individual – a heavy imbalance. By changing the unequalness in the dataset, we are able to investigate if eSVD-DE requires strong assumptions on how evenly distributed the cells are distributed across individuals to maintain high power. (Mechanically, for the last setting, to generate the unequal distribution of cells per individual, we set the number of cells per individual to vary from $250/4 \approx 63$ to $250 \times 4 = 1000$ equally-spaced across the 10 case individuals, and similarly for the 10 control individuals. This yields the aforementioned maximum 16x difference between any two individuals.)

This gives us a total of 9 simulation settings, where we pick one of three number of genes and one of three unequalness of number of cells per individual.

#### C.3 Results of extended power simulations

**ROC curves.** We plot the results as ROC curves in Figure C4. While we had run 25 different trials for each of the 9 different simulation settings to verify that the results were consistent across simulation trials, we only plot the ROC curve from the first trial across the 9 different simulation settings for visual clarity. We verified that the ROC curve shown here is representative of the 24 other ROC curves not shown in this plot.

Interpreting the results, we can notice some apparent trends:

- eSVD-DE performs well across all simulation settings, ranking often as the top method in terms of AUC, as there are many genes in the analysis. (In this case, this is defined as 700 or more genes.) This is sensible, as if there are not many genes, eSVD-DE can not reliably estimating a low-dimensional embedding, in which case it would outperform methods that do not do any embedding.
- SCTransforms performs well in settings where there is heavy imbalance in the number of cells across individuals. This is sensible since as the dataset becomes more imbalanced, it has a less apparent "cohort-level" structure, as most of the cells originate from small percentage of the individuals. In the extreme where most of the cells originate from one case individual and one control individual, the differential expression analysis simplifies to a typical scRNA-seq analysis without any cohort structure.
- MAST performs well in settings when there is an equal distribution of cells across individuals. This is sensible, as MAST uses a mixed effect model with a random intercept per individual. This type of modeling framework is advantageous when each individual has similar number of cells, as this allows the statistical method to accurately estimate the distribution of random intercepts.
- DESeq2 is often a safe alternative. It does not perform poorly in any simulation setting. This further demonstrates that overall, collapsing scRNA-seq data into "pseudobulk" data can yield reasonable results. However, as our simulation demonstrates, because analyzing scRNA-seq on the pseudobulk level necessarily does not take full advantage of the cohort-level scRNA-seq data, resulting in less power compared to eSVD-DE.

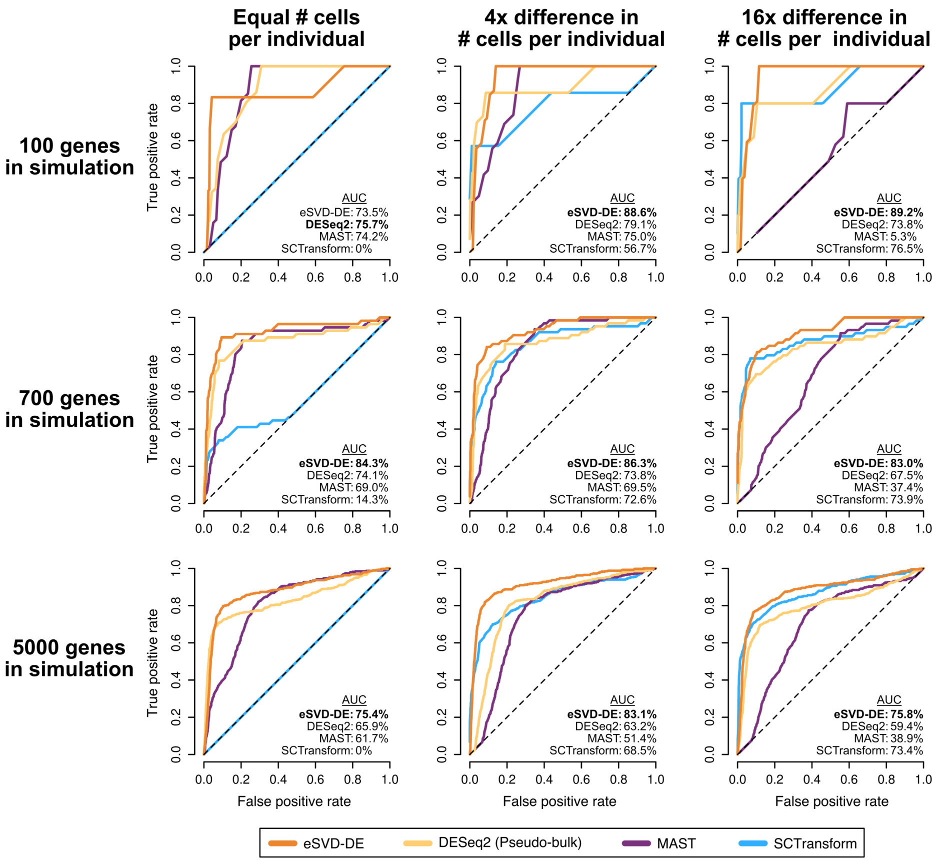


**Fig. C4**: Each ROC curve is similar to Figure 2G, where across the 9 different simulation settings, the ROC curve for the four different methods is shown. The area under the curve (AUC) is shown for each method, where the bolded method denotes the method with the highest AUC. Note that ROC curve for "700 genes" and "Equal # cells per individual" is what is shown in Figure 2G.

**True log-fold change among genes.** For the same simulation datasets across the 9 simulation settings shown in Figure C4, we plot the true log-fold change (defined as the difference in the logarithm of the mean expression among cells from case individuals (unimpacted by the confounders) and the logarithm of the mean expression among cells from control individuals (unimpacted by the confounders) in Figure C5. This is help let the reader gauge the amount of signal in our simulated datasets.

As mentioned in the "Details of power simulation shown in main text" section, most genes do not have a true log-fold change of 0. As discussed there as well, there are five different "clusters" of true log-fold changes due to our assignment of genes to be "strong-negative," "weak-negative," "none," "weak-positive," and "strong-positive." This is to generate synthetic data more reflective of cohort-level scRNA-seq datasets. The genes marked in the red histogram in Figure C5 are the genes deemed "true DE genes" based on the calculating the FDR of each gene based on the true variance and true nuisance parameters and selecting genes with a true FDR less than 0.05.

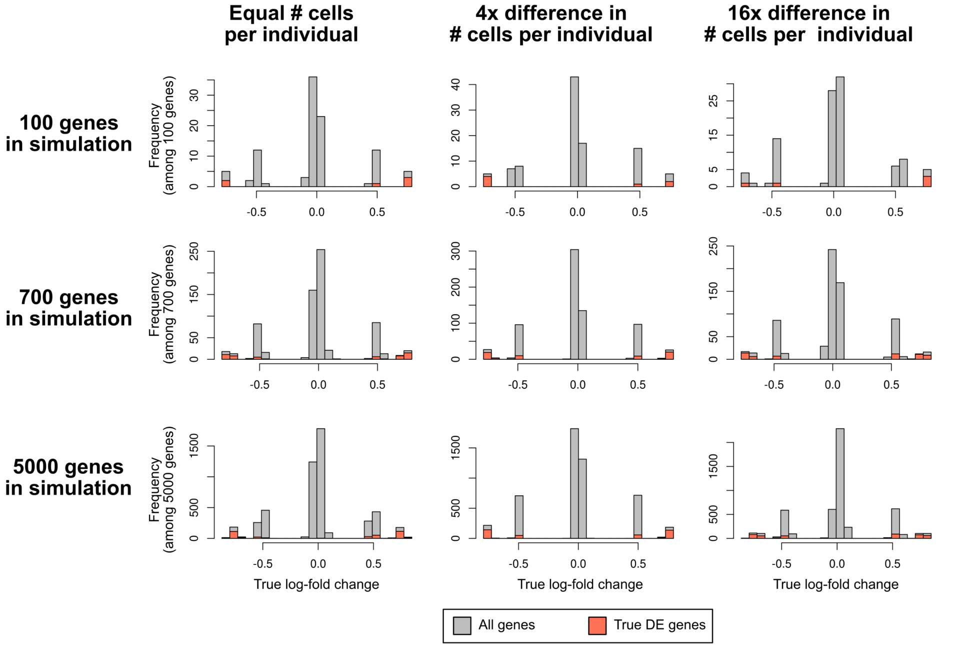


**Fig. C5**: The same simulation settings as shown in Figure C4, where we mark the true log-fold change of each gene in each setting using the (true and unobserved) simulation parameters. The gray histogram marks the frequency of the true log-fold change across all genes, while the red histogram marks the frequency of the true log-fold change across only the "true DE genes" defined based the FDR using the true variance and true nuisance parameters. (The counts in the red histogram is less than or equal to the counts in the gray histogram.)

### Appendix D Computing time and memory usage

We report the computing time and memory usage for each method. Each method was run on one core (i.e., not parallelized) on a Linux server with x86_64 architecture, specifically model: Intel(R) Xeon(R) Platinum 8375C CPU @ 2.90GHz.

**For simulated data.** We first report the computing time and maximum memory usage for the each method when applied on the largest simulated data in the power analysis. The following table is averaged across 10 different trials. This dataset has 5000 genes and 10,626 cells across 20 individuals. Note, the raw simulated data file itself (containing also metadata not directly used in the analysis) has a median size of 0.95 Gigabytes across the 10 trials.

| Method | CPU time (in Hour:Minutes) | Max. Memory (in Gigabytes) |
|---|---|---|
| eSVD-DE | 1:06 | 5.3 |
| DESeq2 | 0:01 | 1.8 |
| MAST | 8:37 | 5.5 |
| SCTransform | 0:02 | 6.9 |

**For Velmeshev dataset (Layer 2/3).** Next, we report the computing time and maximum memory usage for the each method when applied on the Velmeshev dataset shown in Figure 5. This dataset has 7055 genes and 12,984 cells across 31 individuals. Note, the data itself (containing also metadata not directly used in the analysis) has size of 0.2 Gigabytes.

| Method | CPU time (in Hour:Minutes) | Max. Memory (in Gigabytes) |
|---|---|---|
| eSVD-DE | 0:51 | 8.5 |
| DESeq2 | 0:01 | 0.5 |
| MAST | 84:48 | 5.9 |
| SCTransform | 0:08 | 8.1 |

In general, across both the simulated and Velmeshev layer 2/3 data analysis, DESeq2 and SCTransform are very quick to complete (i.e., on the order of minutes), whereas eSVD-DE takes a modest time (i.e., on the order of an hour), and MAST takes a long time to complete, sometimes more than a day. We seek to improve eSVD-DE's optimization sub-routine in future work. MAST's long time to complete stems from its inference of linear mixed models. In terms of memory usage, DESeq2 takes minimal memory due to analyzing pseudobulk data. Otherwise, eSVD-DE, MAST, and SCTransform use similar amounts of memory. While it's hard for us to speculate where the high memory usage stems from for MAST and SCTransform, as we are not familiar with the inner code structure, for eSVD-DE, the high memory usage stems from two reasons: 1) we store dense matrices for the low-dimensional embeddings, and and 2) our implementation stores the initialization as well as the estimated embedding at different rounds (see "Optimization of the embeddings" in the main text). The first point is a fundamental limitation of any low-dimensional embedding method (which we perceive as necessary if we wanted to statistically pool information among cells and genes), and the second point is currently used for ease-of-use when we perform diagnostics on our final estimate. We do not think there are obvious ways to drastically reduce the memory usage of eSVD-DE that is fundamental to the estimation process itself beyond these two points.

### Appendix E Additional details of data preprocessing and analysis

#### E.1 Details of data preprocessing

We expand upon the "Information about data preprocessing" section from the main text. The main aspects we detail here are about: 1) how we select the genes in our analysis, 2) which covariates we include in the analysis, and 3) the parameters we choose when applying the eSVD-DE.

- **Adams et al., and Habermann et al.**: We select T cells for our analysis (i.e., `Manuscript_Identity=="T"`) and the individuals either in the control set or with IPF (i.e., `Disease_Identity %in% c("Control", "IPF")`), and preprocess the scRNA-seq data using `Seurat::NormalizeData` (with `normalization.method`

= "LogNormalize"), Seurat::FindVariableFeatures (with selection.method = "vst" and nfeatures = 5000).

We select T cells for our analysis (i.e., celltype=="T Cells") and the individuals either in the control set or with IPF (i.e., Diagnosis %in% c("Control", "IPF")). To ensure that the analysis between both datasets is comparable, we set the highly variable genes in both datasets to be the 5000 highly variable genes (HVG) in either the Adams and Habermann dataset (after using Seurat::NormalizeData with normalization.method = "LogNormalize" as well as Seurat::FindVariableFeatures with selection.method = "vst" and nfeatures = 5000), as well as the DE genes found for T cells in either [3] or [6], housekeeping genes and cell-cycling genes (from the Seurat package, in cc.genes$s.genes and cc.genes$g2m.genes). This resulted in the 8909 genes used in our analysis for each dataset. Importantly, the usage of Seurat::NormalizeData is soley for defining the highly variable genes, and the eSVD-DE is applied to the original count data.

For analyzing the Adams dataset with eSVD-DE, we use the covariates "Disease_Identity" (which dictate the case-control status), "Subject_Identity" (which dictate which cells originated from which individual), "Gender", "Tobacco", "percent.mt" and "Age". When initializing the eSVD-DE fit, we use $k = 15$ latent dimensional embedding for $X$ and $Y$.

For analyzing the Adams dataset with eSVD-DE, we use the covariates "Diagnosis" (which dictate the case-control status), "Sample_Name" (which dictate which cells originated from which individual), "Gender", "Tobacco", "percent.mt" and "Age". When initializing the eSVD-DE fit, we use $k = 15$ latent dimensional embedding for $X$ and $Y$, and

- **Smillie et al.**: We focus on the cell types encoded in Celltype to be "Cycling TA", "Enterocyte Progenitors", "TA 1", or "TA 2". We remove cells that have one of the highest gene expressions in too many genes, as well as genes that are too sparsely observed. To select the genes for our analysis, we use the highly variable genes (after using Seurat::NormalizeData with normalization.method = "LogNormalize" as well as Seurat::FindVariableFeatures (with selection.method = "vst" and nfeatures = 5000), as well as any DE genes found by [7] (when comparing inflamed to healthy, non-inflamed to healthy, or inflamed to non-inflamed), the housekeeping genes, and the cell-cycling genes. Importantly, the usage of Seurat::NormalizeData is soley for defining the highly variable genes, and the eSVD-DE is applied to the original count data. Finally, we keep the cells whose variance across all the selected genes is not too low.

  When analyzing any cell type using eSVD-DE, we first compare the healthy individuals (i.e., Subject_Disease == "HC" and Sample_Health == "Healthy") to case individuals with a tissue section that is inflamed (i.e., Subject_Disease == "Colitis" and Sample_Health == "Inflamed"). We perform a separate analysis that compares the healthy individuals to case individuals with a tissue section that is non-inflamed (i.e., Subject_Disease == "Colitis" and Sample_Health ==

"Non-inflamed"). To ensure that our analysis of healthy vs. inflamed compared to our analysis of healthy vs. non-inflamed shows biologically meaningful correlation, we split the healthy individuals into two groups. We use only one group of healthy individuals in each analysis.

For analyzing the any cell type, we use the covariates "Subject_Disease" (which dictate the case-control status), "Sample" (which dictate which cells originated from which individual), "Subject_Gender", "Subject_Smoking", "Subject_Location", and "percent.mt". When initializing the eSVD-DE fit, we use $k = 15$ latent dimensional embedding for $X$ and $Y$.

- **Velmeshev et al.**: We focus on the cell types encoded in celltype to be "AST-PP", "Endothelial", "IN-SST", "IN-VIP", "L2/3", "L4", "L5/6", "L5/6-CC", "Microglia", "Oligodendrocytes", and "OPC". We remove cells that have one of the highest gene expressions in too many genes, as well as genes that are too sparsely observed. To select the genes for our analysis, we use the highly variable genes (after using `Seurat::NormalizeData` with `normalization.method = "LogNormalize"` as well as `Seurat::FindVariableFeatures` (with `selection.method = "vst"` and `nfeatures = 5000`), as well as any DE genes found by [5], the SFARI genes, the housekeeping genes, and the cell-cycling genes. Importantly, the usage of `Seurat::NormalizeData` is soley for defining the highly variable genes, and the eSVD-DE is applied to the original count data. Finally, we keep the cells whose variance across all the selected genes is not too low.

  For analyzing the any cell type for either of the comparisons, we use the covariates "diagnosis" (which dictate the case-control status), "individual" (which dictate which cells originated from which individual), "region", "age", "sex", "Seqbatch", and "Capbatch". When initializing the eSVD-DE fit, we use $k = 30$ latent dimensional embedding for $X$ and $Y$.

In our application of eSVD-DE, we set the regularization parameter $\tau = 0.1$ for the initialization as well as the alternative minimization.

#### E.2 Tables summarizing all the datasets

In Table E1, we report the summary statistics for each dataset. Here, $p$ is the number of genes, $n$ is the number of cells, $S$ is the number of individuals, and "case" refers to the number of case individuals (strictly smaller than $S$). We also report three key summary statistics of each dataset reflecting its data quality. Here, "cell #" refers to the median number of cells per individual (across all $S$ individuals), "depth" refers to the median sequencing depth of each cell (across all $n$ cells), and "sparsity" refers to proportion of non-zeros in entire $p$-by-$n$ count matrix, as a percentage of the entire matrix (where 100% would refer to a count matrix with no zeros).

**Table E1**: Extended summary table of all datasets

| Dataset | $p$ | $n$ | $S$ | case | cell # | depth | sparsity |
|---|---|---|---|---|---|---|---|
| Adams | 6969 | 8909 | 34 | 24 | 196 | 2562 | 7.1% |
| Habermann | 6969 | 5286 | 10 | 6 | 314 | 3444 | 8.4% |
| Smillie: Cycling TA (I) | 6604 | 4856 | 14 | 8 | 126 | 9011 | 17.3% |
| Smillie: Cycling TA (NI) | 6604 | 5788 | 14 | 8 | 382 | 10,328 | 19.9% |
| Smillie: Enter. Prog. (I) | 6013 | 2344 | 10 | 4 | 110 | 1084 | 6.3% |
| Smillie: Enter. Prog. (NI) | 6013 | 3641 | 10 | 4 | 273 | 816 | 5.5% |
| Smillie: TA 1 (I) | 5713 | 6380 | 19 | 13 | 167 | 903 | 4.8% |
| Smillie: TA 1 (NI) | 5713 | 14,215 | 19 | 13 | 477 | 749 | 4.4% |
| Smillie: TA 2 (I) | 6617 | 3055 | 12 | 6 | 129 | 7221 | 15.5% |
| Smillie: TA 2 (NI) | 5713 | 3801 | 12 | 6 | 350 | 9350 | 19% |
| Velmeshev: AST-PP | 5943 | 7559 | 31 | 15 | 218 | 1484 | 14.5% |
| Velmeshev: Endothe. | 6085 | 2600 | 31 | 15 | 67 | 1243 | 14.3% |
| Velmeshev: IN-SST | 6691 | 4411 | 31 | 15 | 131 | 3591 | 21.5% |
| Velmeshev: IN-VIP | 6887 | 5839 | 31 | 15 | 163 | 4028 | 21.8% |
| Velmeshev: L2/3 | 7094 | 12,984 | 31 | 15 | 326 | 8799 | 33.7% |
| Velmeshev: L4 | 7041 | 6666 | 31 | 15 | 167 | 6293 | 28% |
| Velmeshev: L5/6 | 7087 | 3459 | 31 | 15 | 107 | 7699 | 30.4% |
| Velmeshev: L5/6-CC | 7146 | 4484 | 31 | 15 | 124 | 12,499 | 38.9% |
| Velmeshev: Microglia | 4079 | 3320 | 29 | 15 | 102 | 616 | 12.5% |
| Velmeshev: Oligo | 4611 | 15,194 | 31 | 15 | 306 | 896 | 14.4% |
| Velmeshev: OPC | 5696 | 9954 | 31 | 15 | 276 | 1466 | 15% |

#### E.3 Details of visualizing the embedding

To visualize the embeddings (either of the principal components of the log-normalized scRNA-seq data or of the low-dimensional embedding $X$ estimated by eSVD-DE, throughout Figures 2 through 5), we use the `dimRed::embed` function (with `.method="Isomap"` and `knn=30`).

#### E.4 Details of exploring the relationship between expression and sequencing depth

To explore the relationship between the gene expression and sequencing depth (as demonstrated in Figure 3), we deploy a procedure similar to the one used in [8].

Specifically, for the "Before eSVD-DE" analysis, we first cluster all the genes into 6 clusters based on the logarithm of the total expression across all cells. Then, for each gene separately, we regress the gene's expression (across all the cells) onto the log sequencing depth (across all the cells) via a kernel nonparametric regression using the `npregfast::frfast` function. After performing this regression, we compute the predicted gene expression for different log sequencing depths. Finally, for all the genes within one of the 6 clusters, we compute the median as well as the upper and lower quantiles for the predicted gene expression for different log sequencing depths. This is visualized in Figure 3C.

For the "After eSVD-DE" analysis, we apply the same procedure, except after fitting the eSVD-DE, we regress the denoised gene's expression, not including the covariate-adjusted sequencing depth (across all $n$ cells), i.e., for gene $j$,

$$\left\{\hat{\mu}_{ji}\right\}_{i=1}^{n} = \left\{\exp\left((\hat{Y}_{j,\cdot})^\top(\hat{X}_{i,\cdot}) + \hat{Z}_{j,(\mathrm{cc})} \cdot C_{i,(\mathrm{cc})}\right)\right\}_{i=1}^{n},$$

onto the log sequencing depth (across all the cells) via a nonparametric regression, and then proceed with the remainder of the analysis similar to the "Before eSVD-DE" analysis. This is visualized in Figure 3D.

#### E.5 Details of DESeq2

For all analyses using DESeq2 [9], we first construct pseudobulk samples for each individual, whereby for each gene, we sum the counts across all the cells from that individual. If there is no variation among a covariate among all the cells for this individual, we set the covariate for the pseudobulk to be equal to the unique covariate from the individual. We use the mean value for covariates such as `percent.mt` that vary across the cells. Then, we use `DESeq2::DESeqDataSetFromMatrix` and `DESeq2::DESeq` to perform the DE test, whereby we use a Benjamini-Hochberg multiple-testing correction afterward.

#### E.6 Details of MAST

We first log-normalize the RNA counts for all analyses using MAST [10]. Then, we use the `MAST::FromMatrix` function (setting `cData` accordingly based on the cells' covariates) and the `MAST::zlm` function to fit the mixed-effects model, using a random effect for the individual (with `method = "glmer"` and `ebayes = FALSE`), followed by the `MAST::lrTest` function to perform the DE test, whereby we use a Benjamini-Hochberg multiple-testing correction afterward.

#### E.7 Details of SCTransform

For all analyses using SCTransform [8], we use the `Seurat::SCTransform` (with `method = "glmGamPoi"`) and set the `vars.to.regress` argument to be all the relevant covariates (notably, not including any one-hot encoding vectors that depict which cell originated from which individual). Then, we use `Seurat::FindMarkers` function (with `test.use = "wilcox"`, `logfc.threshold = 0`, and `min.pct = 0`) to perform the DE test, whereby we use a Benjamini-Hochberg multiple-testing correction afterward.

#### E.8 Details of GLM-PCA

For all analyses using GLM-PCA [11], we use the `glmpca::glmpca` (with `fam = "poi"`, `minibatch = "stochastic"`) with an equal number of latent dimensions as the corresponding eSVD-DE analysis, and setting `X` to be the relevant covariates (excluding the case-control covariate) Then, after the extracting the fitted factors and loadings that do not correspond to the effects of the covariates, we construct the matrix $\mu \in \mathbb{R}^{p\times n}$, the estimated mean matrix (i.e., exponentiated natural parameter matrix). We then perform a Wilcoxon rank sum test via `stats::wilcox.test` on each gene to perform the DE test, whereby we use a Benjamini-Hochberg multiple-testing correction afterward.

##### E.9 Details of enrichment analysis

For the GO enrichment analysis in Figure 5, we used the `clusterProfiler::enrichGO` function (with `OrgDb = org.Hs.eg.db::org.Hs.eg.db`, `keyType = "SYMBOL"`, `ont = "BP"`, `pAdjustMethod = "BH"`, `pvalueCutoff = 0.05`, and `qvalueCutoff = 0.05`.

For the treemaps as shown in Figure F12, we additionally take the output of `clusterProfiler::enrichGO` and pass it into `rrvgo::calculateSimMatrix` (with `method="Rel"`), `rrvgo::reduceSimMatrix` (setting the `scores` input to be the negative $\log_{10}$ q-values from `clusterProfiler::enrichGO`, and with `threshold=0.7`), and `rrvgo::treemapPlot`.

#### Appendix F Additional data analysis results

The following figures provide supplementary results of all these analyses mentioned in the main text.

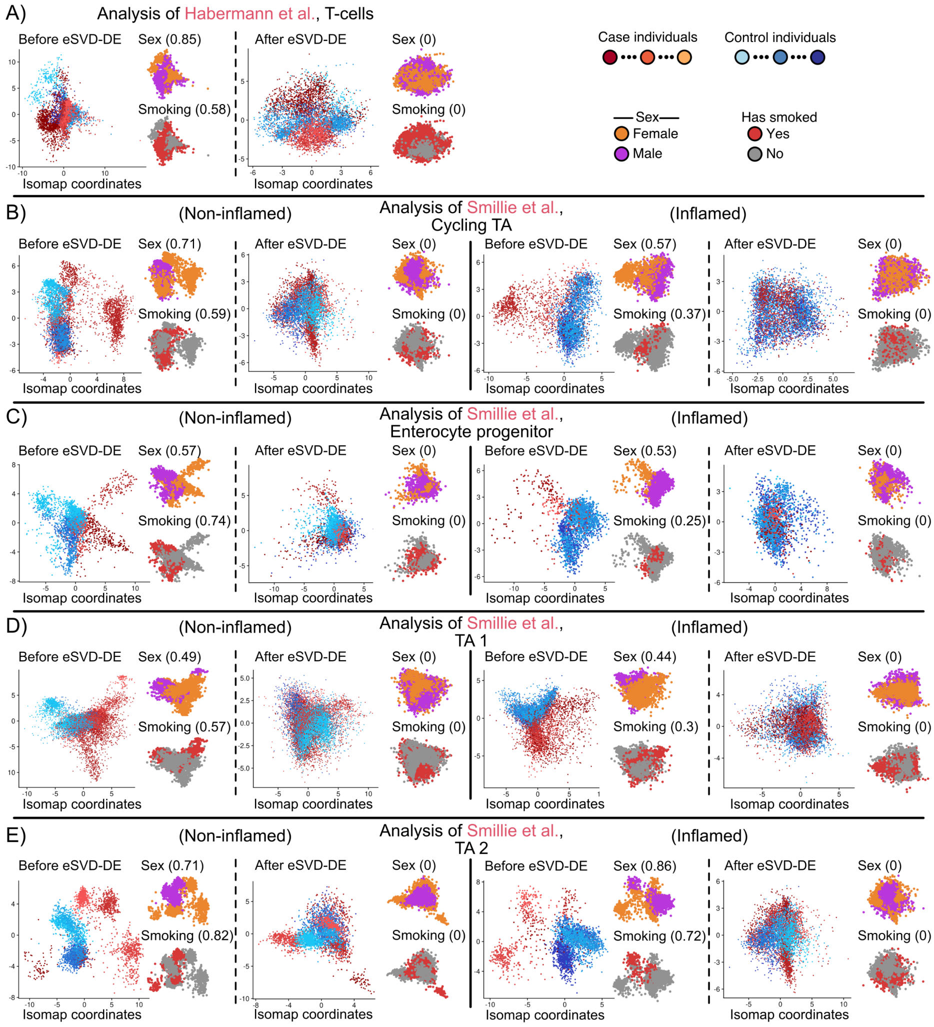


**Fig. F6**: Each Isomap is similar to Figure 3(A,B), but for a different dataset, the Isomap is constructed on the leading principal components (i.e., before eSVD-DE) or based on the low-dimensional cell embedding $X$ after applying the eSVD-DE. We primarily show the Isomaps where the cells are colored by the different case and control individuals that contributed to the cell and also show different insets where the cells are colored by the sex of the individual as well as whether or not the individual smoked. The ideal Isomap after applying eSVD-DE shows no discernable separation between individuals of different sex and smoking status, as these covariates should have been regressed out. **A** Isomaps of the T-cells in the Habermann dataset [6]. **B,C,D,E** Isomaps of the cycling TAs, enterocyte progenitors, TA 1's, or TA 2's in the Habermann dataset [6], either for healthy and non-inflamed cells (left) or healthy and inflamed cells (right).


[6] Habermann, A.C., Gutierrez, A.J., Bui, L.T., Yahn, S.L., Winters, N.I., Calvi, C.L., Peter, L., Chung, M.-I., Taylor, C.J., Jetter, C., Raju, L., Roberson, J., Ding, G., Wood, L., Sucre, J.M.S., Richmond, B.W., Serezani, A.P., McDonnell, W.J., Mallal, S.B., Bacchetta, M.J., Loyd, J.E., Shaver, C.M., Ware, L.B., Bremner, R., Walia, R., Blackwell, T.S., Banovich, N.E., Kropski, J.A.: Single-cell

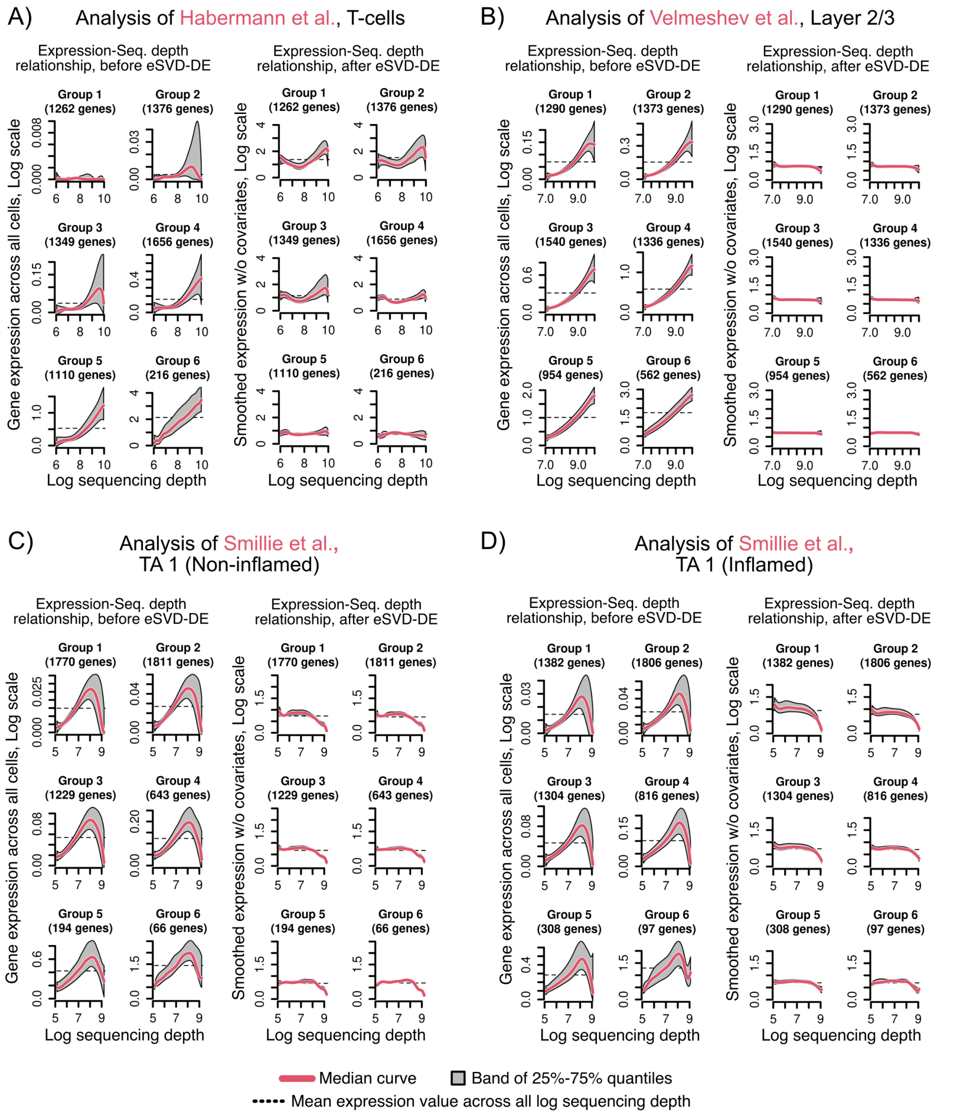


**Fig. F7**: Each line plot is similar to Figure 3(C,D), but for a different dataset, genes are partitioned into six bins based on the gene's mean expression. **A** Relationship between gene expression and sequencing depth for the T-cells in the Habermann dataset [6] before and after applying eSVD-DE. **B** Relationship between gene expression and sequencing depth for the layer 2/3 cells in the Velmeshev dataset [5] before and after applying eSVD-DE. **C,D** Relationship between gene expression and sequencing depth for the TA 1 cells in the Smillie dataset [7] before and after applying eSVD-DE, either for the healthy and inflamed cells (**C**) or the healthy and non-inflamed cells (**D**). An ideal relationship after applying eSVD-DE is that the gene expression is constant for different sequencing depths for each bin of genes.

RNA sequencing reveals profibrotic roles of distinct epithelial and mesenchymal lineages in pulmonary fibrosis. Science Advances **6**(28), 1972 (2020)

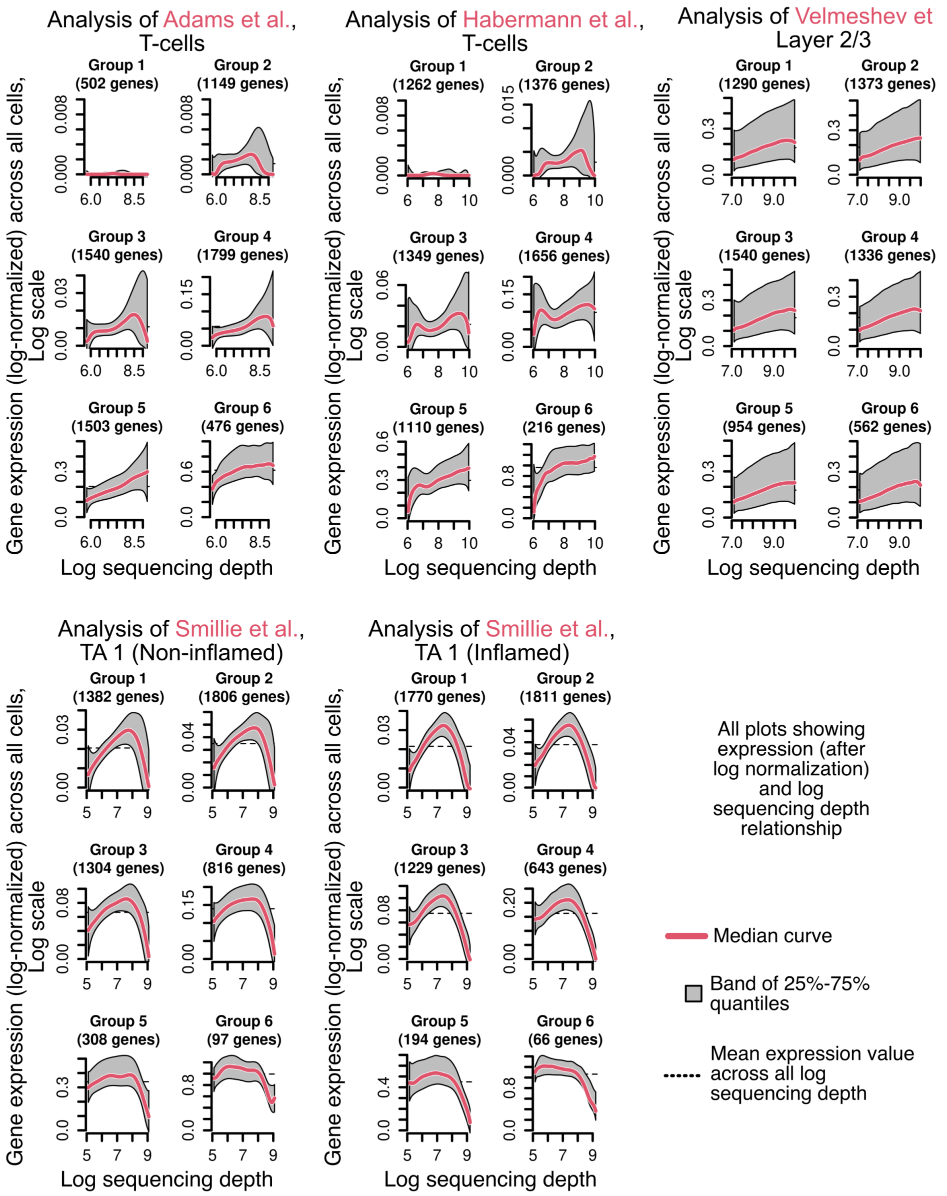


**Fig. F8**: Analogous plots to Figure 3(C,D) and F7, but shown if the gene expressions in each scRNA-seq dataset were log-normalized instead using eSVD-DE. The Y axis denotes the log-normalized gene expression across the same gene partitions as in Figure 3(C,D) and F7 (on an additionally applied log scale). This latter application of the log scale is to keep the plots here as comparable to those in Figure 3(C,D) and F7 as possible. We see that in all the five datasets, log-normalization does not fully mitigate the relationships between gene expression and sequencing depth. This means after log-normalization, genes might still be assess to be differentially expressed solely based on differences in sequencing depth.


[7] Smillie, C.S., Biton, M., Ordovas-Montanes, J., Sullivan, K.M., Burgin, G., Graham, D.B., Herbst, R.H., Rogel, N., Slyper, M., Waldman, J., Sud, M., Andrews,

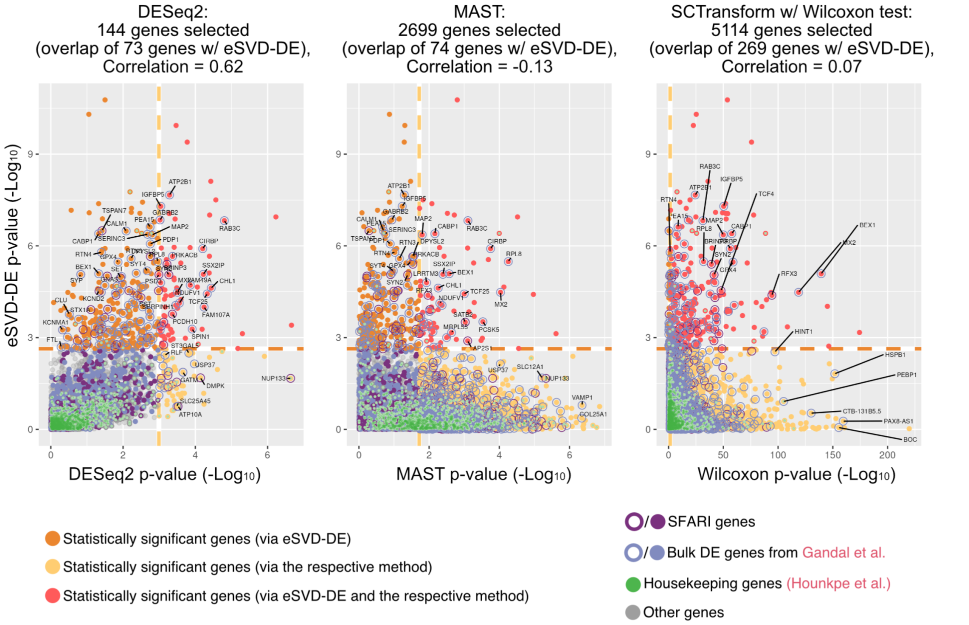


**Fig. F9**: Scatter plots comparing the negative $\log_{10}$ p-values of all 7055 genes derived via eSVD-DE against either DESeq2, MAST, or SCTransform (followed by the Wilcoxon rank-sum test). Genes with a Benjamini-Hochberg multiple-testing corrected p-value less than 0.05 are deemed significant within their respective method. The number of significant genes for each of the three methods is written in the title, alongside the number of genes that overlap with eSVD-DE's 331 significant genes. The correlation between both sets of negative $\log_{10}$ p-values is also noted. Note that both MAST and SCTransform deem a large proportion of the 7055 genes to be statistically significant.

E., Velonias, G., Haber, A.L., Jagadeesh, K., Vickovic, S., Yao, J., Stevens, C., Dionne, D., Nguyen, L.T., Villani, A.-C., Hofree, M., Creasey, E.A., Huang, H., Rozenblatt-Rosen, O., Garber, J.J., Khalili, H., Desch, A.N., Daly, M.J., Ananthakrishnan, A.N., Shalek, A.K., Xavier, R.J., Regev, A.: Intra-and inter-cellular rewiring of the human colon during ulcerative colitis. Cell **178**(3), 714–730 (2019)

[8] Hafemeister, C., Satija, R.: Normalization and variance stabilization of single-cell RNA-seq data using regularized negative binomial regression. Genome Biology **20**(1), 1–15 (2019)

[9] Love, M.I., Huber, W., Anders, S.: Moderated estimation of fold change and dispersion for RNA-seq data with DESeq2. Genome Biology **15**(12), 550 (2014)

[10] Finak, G., McDavid, A., Yajima, M., Deng, J., Gersuk, V., Shalek, A.K., Slichter, C.K., Miller, H.W., McElrath, M.J., Prlic, M., Linsley, P.S., Gottardo, R.: MAST: A flexible statistical framework for assessing transcriptional changes and characterizing heterogeneity in single-cell RNA sequencing data. Genome Biology **16**(1), 278 (2015)

[11] Townes, F.W., Hicks, S.C., Aryee, M.J., Irizarry, R.A.: Feature selection and

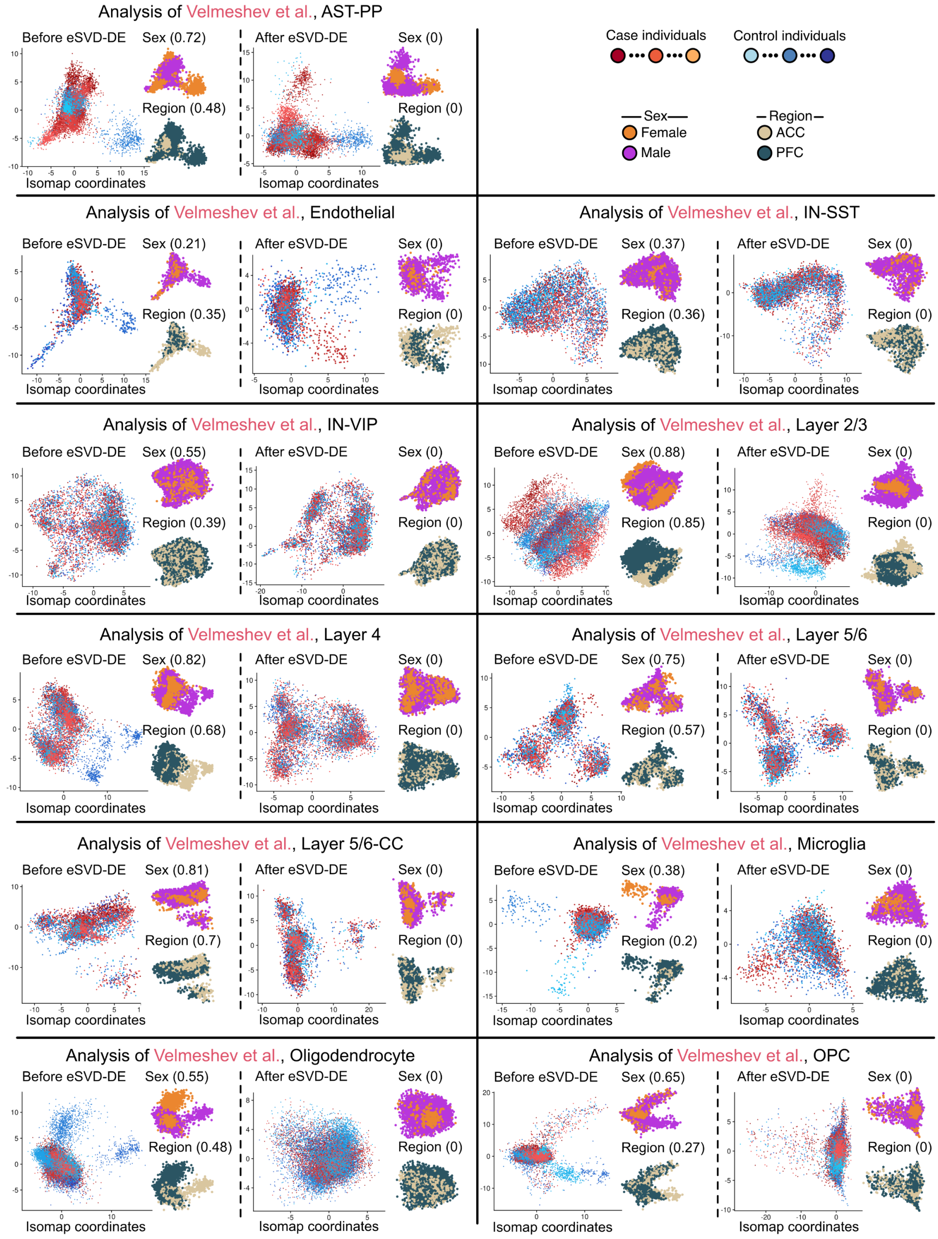


**Fig. F10**: A continuation of Figure F6, where the different rows show the Isomaps for different cell types in the Velmeshev dataset [5], where for any cell type, the left plot shows the Isomap on the leading principal components (i.e., before applying eSVD-DE), and the right plot shows the low-dimensional space $X$ after applying eSVD-DE.

dimension reduction for single-cell RNA-seq based on a multinomial model. Genome Biology **20**(1), 1–16 (2019)

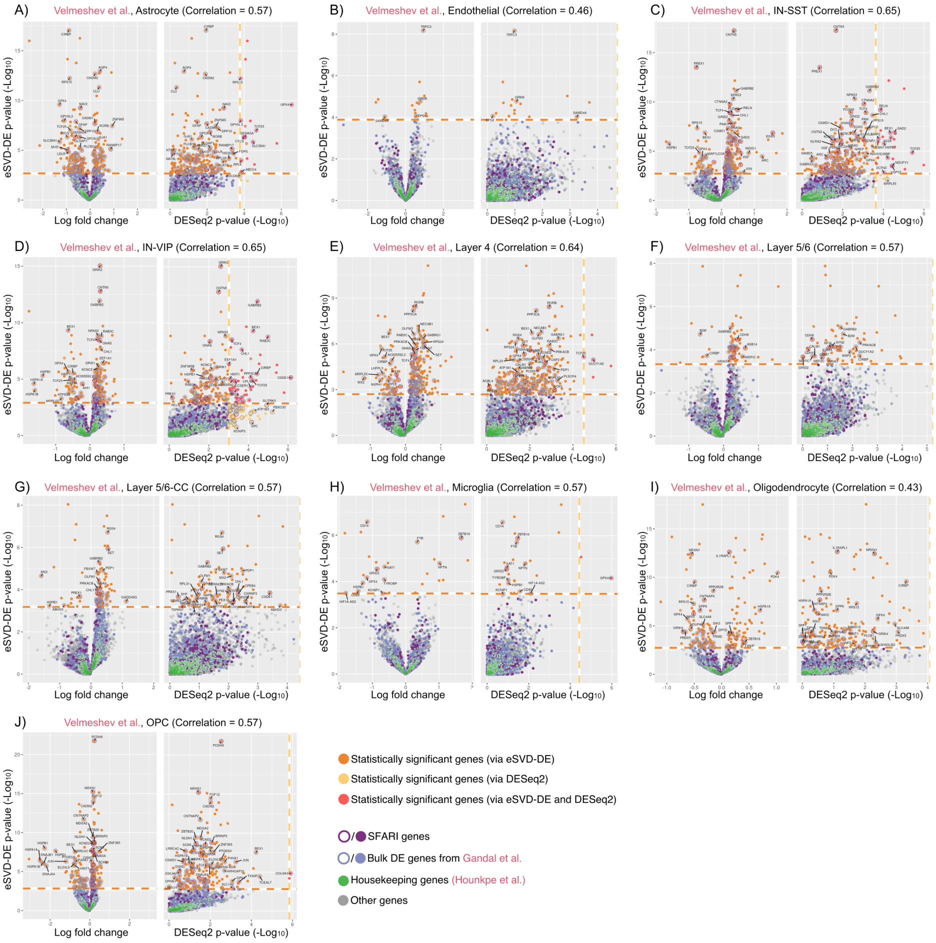


**Fig. F11**: Each volcano plot is similar to Figure 5(B,C), but applied to a different cell type in the Velmeshev dataset [5]. Each gene in every plot is marked differently, depending on whether or not it was deemed significant by eSVD-DE or DESeq2, as well as if the gene was a SFARI gene, a bulk DE gene, or housekeepin gene. The correlation written in the title of each plot denotes the correlation between the negative $\log_{10}$ p-values between the eSVD-DE results and the DESeq2 results. **A** The astrocytes. **B** The endothelial cells. **C** The IN-SST cells. **D** The IN-VIP cells. **E** The layer 4 cells. **F** The layer 5/6 cells. **G** The layer 5/6-cc cells. **H** The microglias **I** The oligodendrocytes. **J** The OPCs.

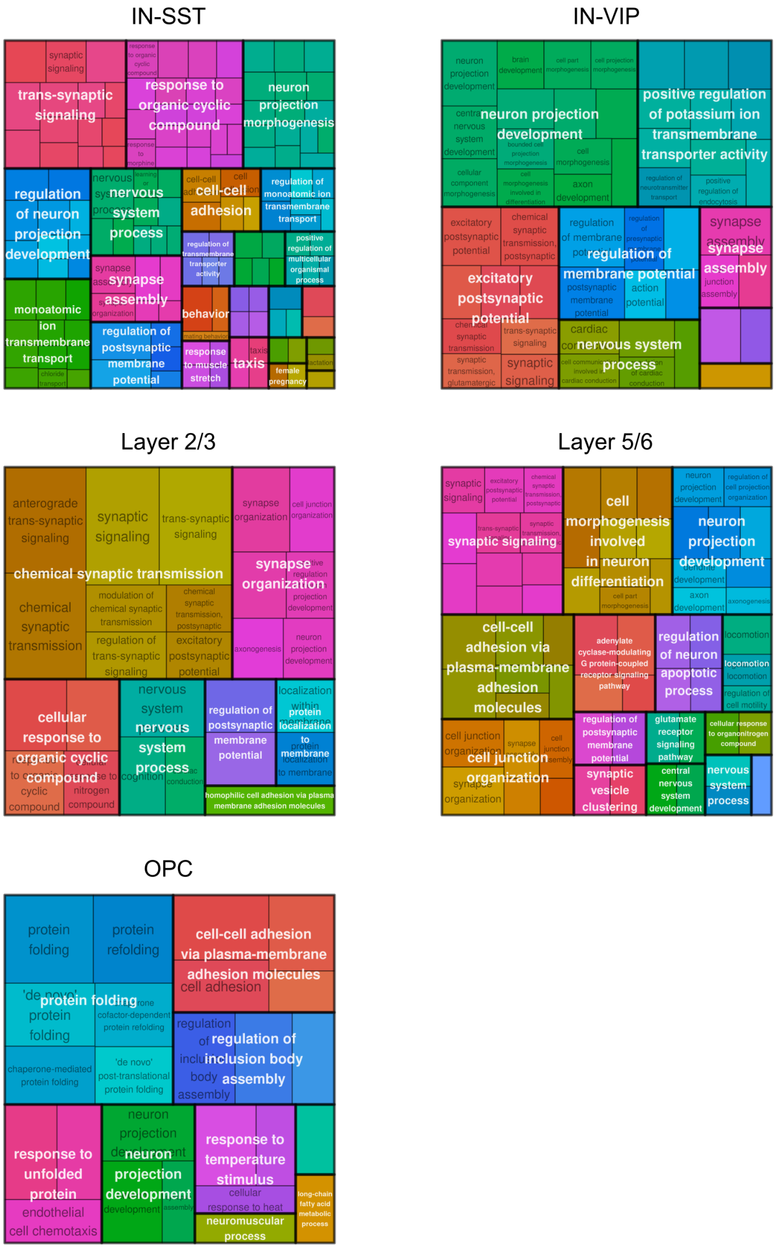


**Fig. F12**: Tree maps of the enriched GO terms among the significant genes discovered by eSVD-DE when analyzing different cell types. While this analysis was run on every cell type, some significant genes were not enriched enough GO terms to form a tree map (and hence, not shown among the five shown in this plot). Each plot shows different enriched GO terms (deemed significant by Fisher's exact test p-value after Benjamini-Hochberg multiple-testing correction), grouped by two hierarchies, where the top hierarchy is shaded with a different color.
